# A Robust Fluorogenic Substrate for Chikungunya Virus Protease (nsP2) Activity

**DOI:** 10.1101/2025.02.01.636040

**Authors:** Sparsh Makhaik, Wioletta Rut, Shruti Choudhary, Tulsi Upadhyay, Chenzhou Hao, Michael Westberg, Cedric Bobst, Euna Yoo, Jasna Fejzo, Michael Z Lin, Matthew Bogyo, Paul Thompson, Marcin Drag, Jeanne A. Hardy

## Abstract

Chikungunya virus (CHIKV) is an emerging pathogen with pandemic potential. CHIKV infection in humans is transmitted by mosquitoes and induces common symptoms of high fever, arthralgia and myalgia. Because no specific antiviral drugs for treatment of CHIKV infection are available, drug development remains a central goal. The chikungunya virus protease nsP2 (CHIKVP) has emerged as a key drug target due to its indispensable role in viral replication via cleavage of the viral polyprotein. To date, effective tools for screening for CHIKVP inhibitors that reflect the most critical polyprotein cleavage sites have been lacking, hampering drug-development efforts. We found that the recognition ability of CHIKVP is sensitive to the length of peptide substrates. In this study, we report a robust fluorogenic substrate comprising a 15-mer peptide derived from the nsP3/4 junction from the CHIKV polyprotein. This peptide is flanked by an ACC-Lys(dnp) donor-quencher pair. Our new substrate acc-CHIK_15_-dnp shows a 20-fold improved signal-to-noise ratio as compared to the previously reported edab_8_ substrate, which is also based on the nsP3/4 junction. We found acc-CHIK_15_-dnp is recognized only by CHIKVP but not by other alphavirus proteases. This is surprising due to the high level of sequence conservation in the alpha virus polyprotein junctions and indicates that the P-side residues are more important than the P’-side sequence for effective CHIKVP cleavage. The robust signal-to-noise ratio obtained using acc-CHIK_15_-dnp derived from the nsP3/4 cleavage site enabled much improved small molecule HTS on CHIKV relative to other fluorogenic reporters.

## INTRODUCTION

Chikungunya virus (CHIKV) is a crippling arthropod-borne disease of pandemic potential spread by *Aedes aegypti* or *Aedes albopictus* mosquito. CHIKV was first reported in East Africa (Lumsden, 1955; Robinson, 1955) and has now been detected across Asia, Europe and the Americas infecting over 3 million people annually (Chikungunya Virus CDC, 2024). Symptoms of acute CHIKV infection include high fever, headache, rash, and muscle and joint pain (Sadanand, 2018), whereas chronic symptoms may result in debilitating joint swelling as well as incapacitating stiffness and arthralgia mimicking rheumatoid arthritis (Amaral, Bilsborrow, & Schoen, 2020). CHIKV infection during late pregnancy can also result in severe encephalopathy in newborns and aborted fetuses (Gérardin et al., 2014). Fortunately, IXCHIQ^®^, a single-shot vaccine developed by Valneva was recently approved by the FDA (Valneva SE, 2023) which should positively impact infection rates overall. Nevertheless, due to its mounting infection rate and extreme debilitating impact, there is a great need for antiviral drugs for the treatment of CHIKV disease.

CHIKV is an enveloped positive-strand RNA alphavirus and member of the *Togaviridae* family (Schmaljohn AL, 1996). The single-stranded RNA encodes both structural and non-structural proteins (nsPs) (Strauss JH, 1994) of which nsP2 protease (CHIKVP) is a promising drug target due to its role in cleavage of the viral polyprotein. Polyprotein cleavage results in release of five structural proteins: capsid and envelope glycoproteins E1, E2, E3, and 6K (Strauss JH, 1994) and four non-structural proteins: nsP1, nsP2, nsP3 and nsP4 (Strauss JH, 1994) (Figure 1A). nsP2 is a multi-domain protein comprising a nucleoside triphosphatase (Karpe, Aher, & Lole, 2011), RNA dependent RNA helicase (Das, Merits, & Lulla, 2014; Law et al., 2019) and a cysteine protease, CHIKVP. Thus, nsP2 plays essential roles in viral reproduction. CHIKVP contains a cysteine-histidine catalytic dyad (Rausalu et al., 2016), contributing to viral replication by cutting the polyprotein first at nsP3/4 junction (RAGG/YIFS) followed by nsP1/2 (RAGA/GIIE) and lastly separating nsP2/3 (RAGC/APSY) into distinct proteins (Jose J, 2009; A. Lulla, Lulla, & Merits, 2012; A. Lulla, Lulla, Tints, Ahola, & Merits, 2006) (Figure 1B). Notably, these sites all contain one positively charged residue followed by three small residues N-terminal to the cleavage site. Cleavage of the nsP3/4 junction in alphaviruses is fully essential for viral replication, whereas blocking cleavage at the nsP1/2 and nsP2/3 junctions by mutagenesis in Semliki Forest Virus (SFV) (V., Lulla, Karo Astover, Rausalu, Merits, & Lulla, 2013) and Sindbis (Gorchakov et al., 2008; Mai, Sawicki, & Sawicki, 2009) alphaviruses did not prevent viral replication, underscoring the importance of assessing CHIKVP activity at the nsP3/4 junction. Due to its indispensable role in viral duplication, CHIKVP has been identified as a crucial drug target for the development of antivirals against CHIKV. The successful exploitation of viral proteases of HIV (Fox, 1996; William Cameron et al., 1999), Hepatitis-C virus (Foster et al., 2015; Lin, Kwong, & Lin, 2004) and SARS-CoV-2 (Owen et al., 2021) to treat viral infections underscores the relevance of developing protease inhibitors against viral diseases. However, one of the challenges in discovering inhibitors remains development of a robust assay to perform high-throughput screening (HTS) methods on the relevant viral enzyme.

**Figure 1.**
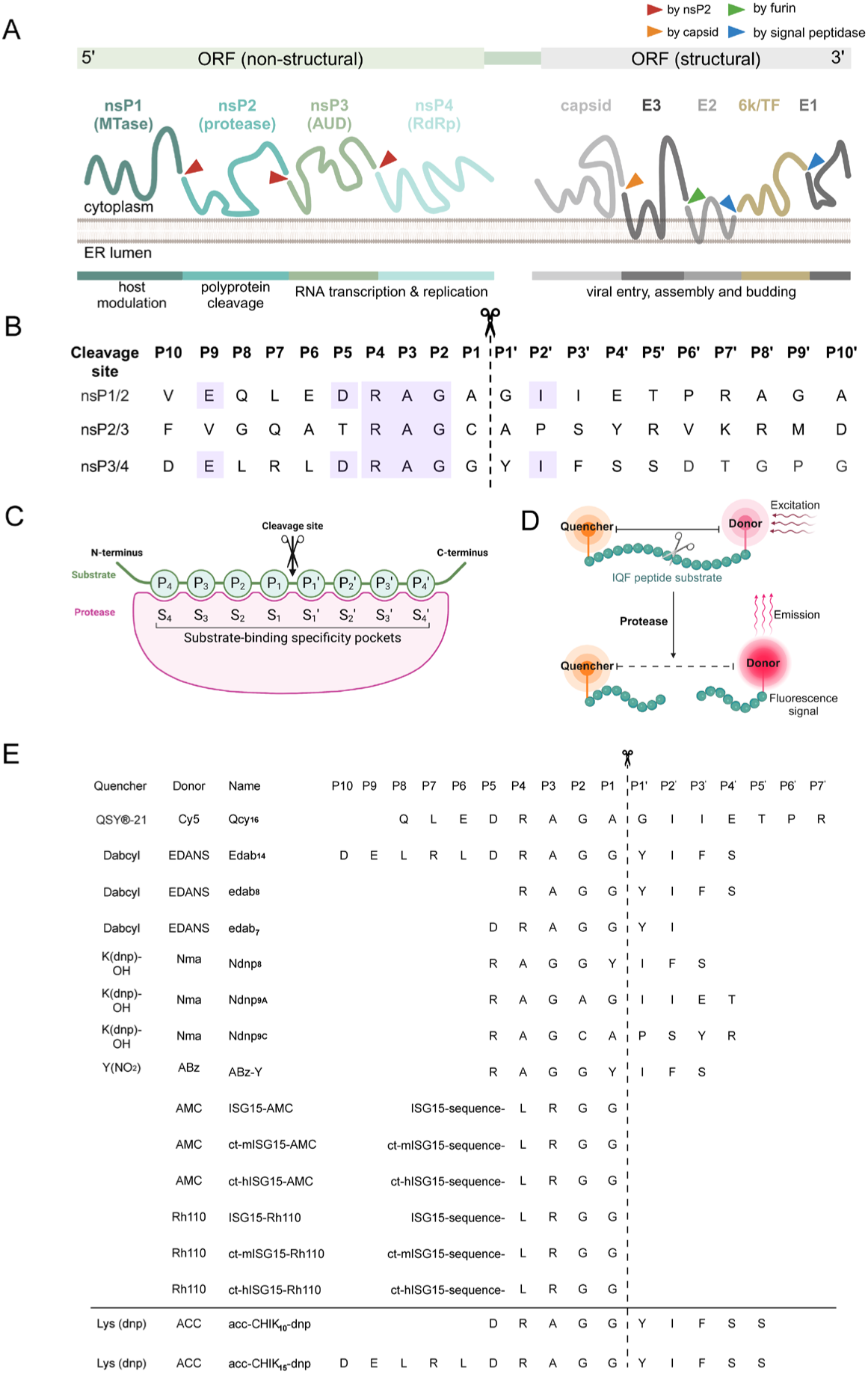
Characteristics of alpha virus polyprotein junctions and fluorogenic substrate derived from polyprotein cleavage sites. **(A)** The polyprotein of CHIKV contains both structural and non-structural proteins. The non-structural ORF encodes non-structural proteins: nsP1 is a methyl transferase (Mtase), nsP2 is a cysteine protease, nsP3 is an alphavirus unique domain (AUD) and nsP4 is the core RNA-dependent RNA polymerase (RdRp). This non-structural ORF is linked via junction to the structural ORF, which encodes for capsid, three envelope proteins: E3, E2, E1 and 6k/TF proteins. **(B)** Cleavage sites within the CHIKV polyprotein at three nsP junctions, which are cleaved by CHIKVP (nsP2) during viral replication. The highlighted regions show the conserved residues within the P and P’ sides. **(C)** Schechter and Berger nomenclature for the substrate specificity of proteases. The subsites (S) on the protease recognize the peptide residues (P) within the substrate. Proteolytic cleavage occurs between P1 and P1′ positions. **(D)** An outline demonstrating the role of internally quenched peptide substrates with a donor and quencher pair on each end, flanking a protease cleavage site. **(E)** Sequences of peptide substrates derived from the CHIKV cleavage junctions, which have been tested for cleavage by CHIKVP, where: EDANS: 5-((2-Aminoethyl) amino) naphthalene-1-

Proteases cleave substrates by hydrolyzing the peptide bonds between two specific amino acids the nomenclature of which is described by the Schechter and Berger (Schechter & Berger, 1967). The subsites (S) on the protease recognize the peptide residues (P) within the substrate. Proteolytic cleavage occurs between P1 and P1′ positions (Figure 1C). Most synthetic peptide substrates for proteases are designed with fluorophores either directly attached to P1-residue or with a donor-quencher pair attached to opposite termini of scissile peptide bond (Yaron, Carmel, & Katchalski-Katzir, 1979). These pairs utilize non-radiative intramolecular energy transfer between the donor and quencher, which occurs due to the overlap of emission spectrum of the donor by the absorption spectrum of the quencher (Figure 1D). These are referred to as FRET (Förster Resonance Energy Transfer) substrates and have been widely used to study properties of enzymes (Chersi, Sezzi, Romano, Evangelista, & Nista, 1990; Florentin, Sassi, & Roques, 1984). Substrate preference analysis of a closely related alphavirus, SFV (De Groot, Hardy, Shirako, & Strauss, 1990; A. Lulla et al., 2012, 2006), showed that the presence of residues on both P and P’ sides was essential for robust cleavage. In contrast, the cleavage requirements for CHIKVP have been less consistent. Some reports have suggested that CHIKVP substrates composed of solely P-side residues are effective (Pastorino et al., 2008), while other reports suggest that both P and P’ residues are required for effective cleavage by CHIKVP (Saisawang, Sillapee, et al., 2015). A few studies suggest that just P4-P4’ residues are sufficient to detect optimal cleavage activity of CHIKVP (Narwal et al., 2018; Saha et al., 2018; Saisawang, Saitornuang, et al., 2015; Saisawang, Sillapee, et al., 2015) while one study claims that peptide P10-P5’ residues are required to be efficiently cleaved by CHIKVP (Rausalu et al., 2016). The reported kinetic parameters measured for CHIKVP using the most common octapeptide edab_8_ (Figure 1E) are k_cat_ of 3.49 × 10^−3^s^−1^ and k_cat_/K_M_ of 2.33 ×10^2^ M^−1^s^−1^ (Singh et al., 2018). A longer substrate edab_14_ has been reported (Rausalu et al., 2016) to have higher values of k_cat_ (4.1 × 10^−2^ s^1^) and k_cat_/K_M_ (1.64 ×10^2^ M^−1^s^−1^) and seems to be the best nsP3/4 substrate reported so far. However, the activity curves shown in that study did not show the expected trend of increasing signal with increasing concentration of CHIKVP, preventing unambiguous interpretation of those prior results. In the past, there have been various reports of in-silico screening for CHIKVP inhibitor development (Ivanova, Rausalu, Ošeka, et al., 2021; Kovacikova & Van Hemert, 2020; Singh et al., 2018; Souza et al., 2023) and one very recent report on CHIKVP inhibitor using a covalent fragment screening-based approach (Merten et al., 2024) performed using a peptide 16-mer substrate derived from nsP1/2 cleavage site of CHIKV with QSY®-21-Cy5 as a FRET pair. The rationale for use of an nsP1/2 peptide sequence for activity assays remains unclear as there have been previous reports (Pastorino et al., 2008; Saisawang, Sillapee, et al., 2015) which have shown that CHIKVP poorly cleaves this junction. Thus, we identified a need to assess the appropriate peptide length and sequences required for optimal cleavage of an nsP3/4 substrate by CHIKVP. In addition, our initial high-throughput screening using a published substrate (Narwal et al., 2018) suffered from poor performance metrics (vide infra). Thus, we were compelled to find or develop a substrate with properties robust enough to enable reliable identification of CHIKVP inhibitors across a broad chemical space.

In the present study, we have analyzed three peptide sequences of different lengths derived from nsP1/2 and nsP3/4 junctions of CHIKV polyprotein to optimize cleavage. Ultimately, we designed acc-CHIK_15_-dnp which includes the 15 amino acids spanning over P10-P5’ from nsP3/4 flanked by the ACC-Lys (dnp) donor-acceptor pair. acc-CHIK_15_-dnp shows significantly improved kinetic parameters compared to other reported substrates.

## RESULTS

### Selection of peptide sequence for recognition by CHIKVP

Our ultimate goal for this project is to develop inhibitors of CHIKVP, but we recognized that the goal was reliant on accurate assessment of CHIKVP activity. Although a number of fluorescent substrates have been described (Merten et al., 2024; Narwal et al., 2018; Pastorino et al., 2008; Rausalu et al., 2016; Saha et al., 2018; Saisawang, Saitornuang, et al., 2015; Saisawang, Sillapee, et al., 2015; Singh et al., 2018), in our hands, the existing substrates did not have the sufficient signal-to-noise for robust inhibitor identification. Thus, we undertook an exploration of both reported and potential substrates to assess which are readily cleaved by CHIKVP.

For many proteases a simple fluorogenic substrate composed of a short peptide derived from the P-side sequence, covalently attached to a fluorophore (e.g. 7-Amino-4-methylcoumarin) at the P1-cleavage site serves as an excellent monitor of protease activity. In these substrates, the covalent attachment of the peptide quenches the fluorophore, which can be released upon cleavage leading to up to 50,000-fold greater fluorescence (Graham Knight, 1995). Based on reports of cleavage of CHIKV peptide substrates lacking P’ residues (Pastorino et al., 2008; Saha et al., 2018; Saisawang, Sillapee, et al., 2015), we were curious if a simple fluorogenic substrate with GG sequence before the scissile bond could be cleaved by CHIKVP. Similar to the CHIKVP substrate, ubiquitin and ubiquitin-like modifiers such as NEDD8 and ISG15 (Interferon-stimulated gene 15) contain GG sequences at the P-side of the cleavage site (LRGG). We first tested two commercially available short peptide fluorogenic substrates, ac-LRGG-amc and Boc-AGG-amc. Although these two substrates were the only commercially available substrates that consisted of a GG motif like the nsP3/4 junction, an advantage of these two is that they also allowed us to probe two different peptide lengths and two different N-terminal capping groups. Unfortunately, neither substrate was efficiently cleaved by CHIKVP (Figure S1). We also tested a few ISG15-based substrates with different fluorophores (Figure 1E) against CHIKVP, but observed extremely low activity for these substrates as well (Figure S2).

A second class of fluorogenic protease substrates contain a recognition sequence flanked by a fluorophore quencher pair. Cleavage of the substrate results in separation of the fluorophore from the quencher resulting in an increased fluorescent signal. In general, these types of substrates work well for a variety of endopeptidases but rely on i) development of a peptide of correct sequence and length and ii) optimization of the fluorophore-quencher pair. To date all substrates for alpha viruses have relied on sequences derived from the cleavage sites within the polyprotein. We designed experiments on unmodified peptides to initially assess the optimal composition and length for cleavage. We tested the assessed cleavage of three peptides derived from nsP1/2 and nsP3/4 CHIKV junctions using LC-MS (Figure 2A). In contrast to a prior report (Rausalu et al., 2016), the DRAGG**/**YI peptide was not measurably cleaved by CHIKVP and remained intact even after 1 hour of proteolytic digestion (Figure 2B, S3). Longer peptides spanning VEQLEDRAGA/GIIET (nsP1/2) and DELRLDRAGG/YIFSS (nsP3/4) were effectively cleaved upon CHIKVP addition (Figure 2B). To directly assess which peptide was favored for cleavage by CHIKVP, we performed a competition experiment by mixing both the substrates at equal concentration. After 10 minutes incubation with CHIKVP cleaved products for both the peptides were present (Figure 2C, Figure S4) at similar abundance with a 54.1% decrease in peak area for DELRLDRAGG/YIFFS and 52.7% decrease in peak area for VEQLEDRAGA/GIIET. Given that other alphavirus nsP2 proteases cleave nsP3/4 junctions more efficiently before cleaving other sites during replication (Vasiljeva et al., 2003) we prioritized DELRLDRAGG/YIFFS for a new FRET substrate. We additionally favored the nsP3/4 junction since other alphaviruses are capable of replication even when nsP1/2 and nsP2/3 cleavage sites have been blocked by mutagenesis (V., Lulla et al., 2013). We reasoned that a substrate mirroring the most critical junction would generate a better substrate in the context of inhibitor development.

**Figure 2.**
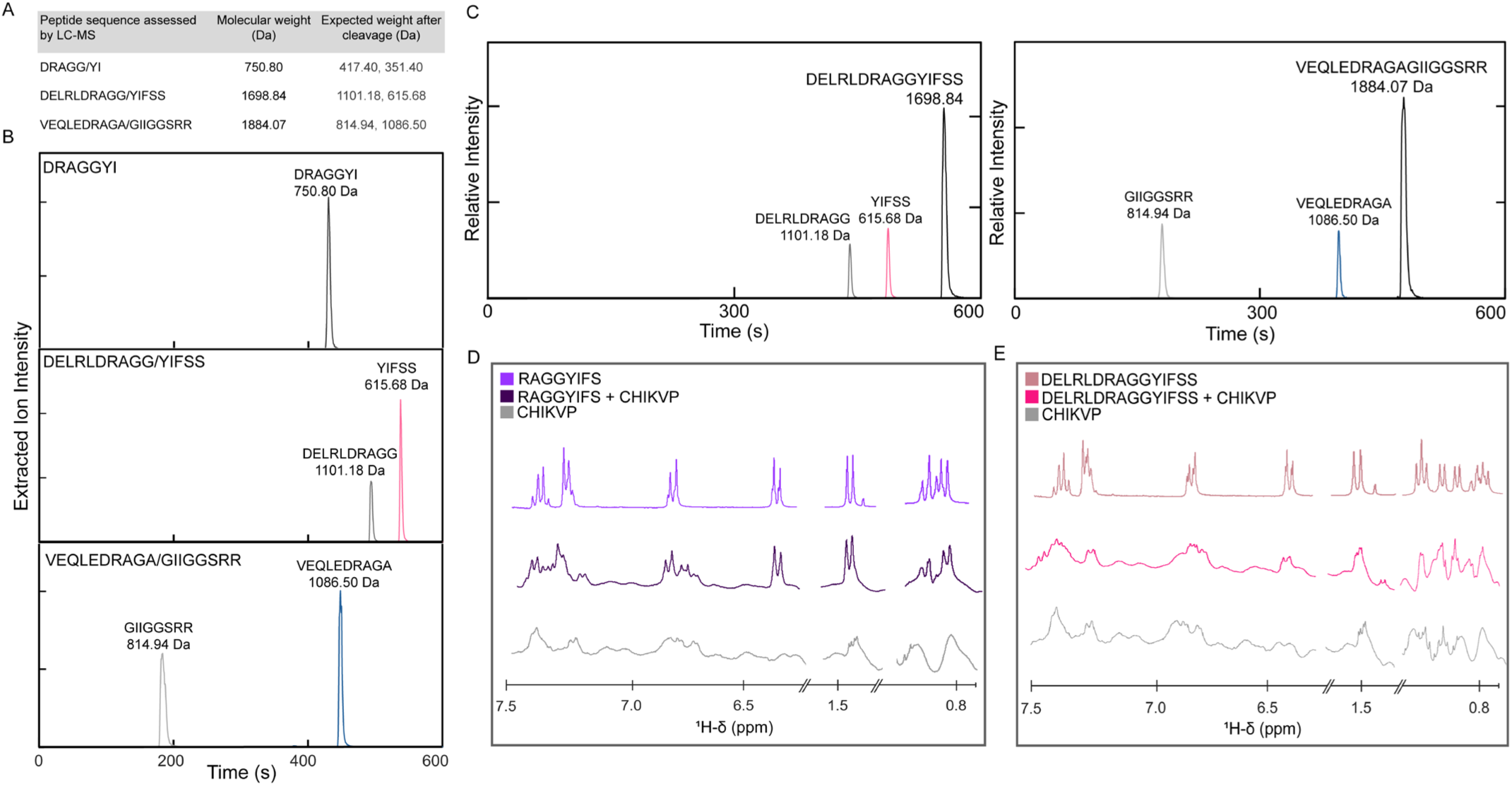
Peptide substrate length impacts CHIKVP recognition. **(A)** Three peptide substrates derived from nsP3/4 or nsP2/3 cleavage sites in CHIKV polyprotein were monitored for cleavage by LC-MS. **(B)** Extracted ion chromatograms show that both of the longer peptides spanning P10 to P5’ residues were cleaved by CHIKVP whereas the substrate lacking residues after P2’ remained uncleaved. **(C)** A CHIKVP cleavage competition assay for two peptides from nsP1/2 and nsP3/4 cleavage sites shows that both are cleaved similarly. **(D)** The substrate RAGGYIFS (residues P4-P4’) was studied by ^1^H-NMR. In the presence of CHIKVP, some line broadening was observed between free and bound peptide suggesting weak binding to CHIKVP. **(E)** The longer substrate DELRLDRAGGYIFSS (residues P10-P5’) showed significant line broadening by ^1^H-NMR suggesting more potent binding to CHIKVP.

Next, we sought insight on binding of the selected substrate as a function of length. The ^1^H-NMR spectra of DELRLDRAGG/YIFSS and RAGG/YIFS in the presence of CHIKVP both showed prominent chemical shifts in the aliphatic (0.5 −1.5 ppm) and aromatic (6.5-7.5 ppm) regions. The chemical shifts from 6.5-7.5 ppm correspond to backbone amide protons and side chain protons. In the presence of CHIKVP significant line broadening of the peptide peaks results from binding to CHIKVP. More significant line broadening is observed for the DELRLDRAGG/YIFSS compared to the RAGG/YIFS (Figure 2D, 2E) suggesting that the longer peptide is a more potent binder of CHIKVP. In addition, the absence of sharp free peptide peaks and massive line broadening observed for DELRLDRAGG/YIFSS suggests that there is chemical exchange (k_ex_) occurring between the free and the bound peptide at the intermediate NMR time scale (Figure S5). However, for RAGG/YIFS the free and the bound peptide exchange fast on the NMR time scale as indicated by less significant line broadening (Figure S5) implying less potent binding. This analysis underscores the selection of the longer peptide DELRLDRAGG/YIFSS derived from the nsP3/4 junction.

### An acc-Lys(dnp) pair results in an effective CHIKVP substrate

The two most commonly used substrates for CHIKVP use EDANS (fluorophore) and Dabcyl (Narwal et al., 2018; Rausalu et al., 2016; Saha et al., 2018; Singh et al., 2018) (quencher) or 2-(N-methylamino)benzoyl (Nma; flourophore) and 2,4-dinitrophenyl (dnp; quencher) (Saisawang, Saitornuang, et al., 2015; Saisawang, Sillapee, et al., 2015) as FRET pairs. A QSY®-21:Cy5 pair was also recently reported (Merten et al., 2024). We observed that the EDANS-Dabcyl and ABz-Y(NO_2_) FRET pairs did not provide a robust readout (Figure S2), so we searched for another fluorophore:quencher pair. These complications have also been reported by others (Matayoshi, Wang, Krafft, & Erickson, 1990; Poreba et al., 2017). The fluorescence of edab_14_ (14-mer CHIKVP substrate with EDANS:Dabcyl derived from nsP3/4 CHIKV site; Figure 1), did not correlate as expected with increasing concentration of CHIKVP (Rausalu et al., 2016). On the other hand, Poreba et. al. reported a highly sensitive and adaptable fluorescence-quenched 7-amino-4-carbamoylmethylcoumarin (ACC) and 2,4-dinitrophenyl-lysine (Lys(dnp)) pair for investigating the substrate specificity of various proteases (Groborz, Kołt, Kasperkiewicz, & Drag, 2019; Kasperkiewicz et al., 2018). The ACC:Lys(dnp) pair shows increased solubility over EDANS:Dabcyl, due to the higher hydrophobicity of Dabcyl relative to the dnp moiety. Dabcyl has also been implicated in undesired interactions with the substrate pockets due to the bulkiness of the Dabcyl moiety (Poreba et al., 2017; Szabó et al., 2021). Moreover, coumarins are considered superior fluorophores to EDANS offering better brightness and therefore enhanced signal for assays (Stewart & Sirinit, 2022). We reasoned that using ACC as a donor and Lys(dnp) as the quencher may be more effective. We designed two new fluorescent substrates based on 10 (P5-P5’) or 15 (P10-P5’) residues from the nsP3/4 cleavage site: acc-CHIK_15_-dnp and acc-CHIK_10_-dnp (Figure 3A, 3B). These substrates showed promising catalytic parameters for monitoring CHIKVP activity - K_M_ of 136.6 µM and k_cat_ of 34.6×10^−2^ s^−1^ for acc-CHIK_15_-dnp and K_M_ of 179.9 µM and k_cat_ of 2.78×10^−2^ s^−1^ for acc-CHIK_10_-dnp. The k_cat_/K_M_ for acc-CHIK_15_-dnp of 2.40×10^3^ M^−1^s^−1^ and 1.54×10^2^ M^−1^s^−1^ for acc-CHIK_10_-dnp suggest that the longer substrate was cleaved more efficiently than the shorter one (Figure 3C). This also suggests the importance of P -side residues over P’-side residues for efficient recognition and cleavage of substrate by CHIKVP.

**Figure 3.**
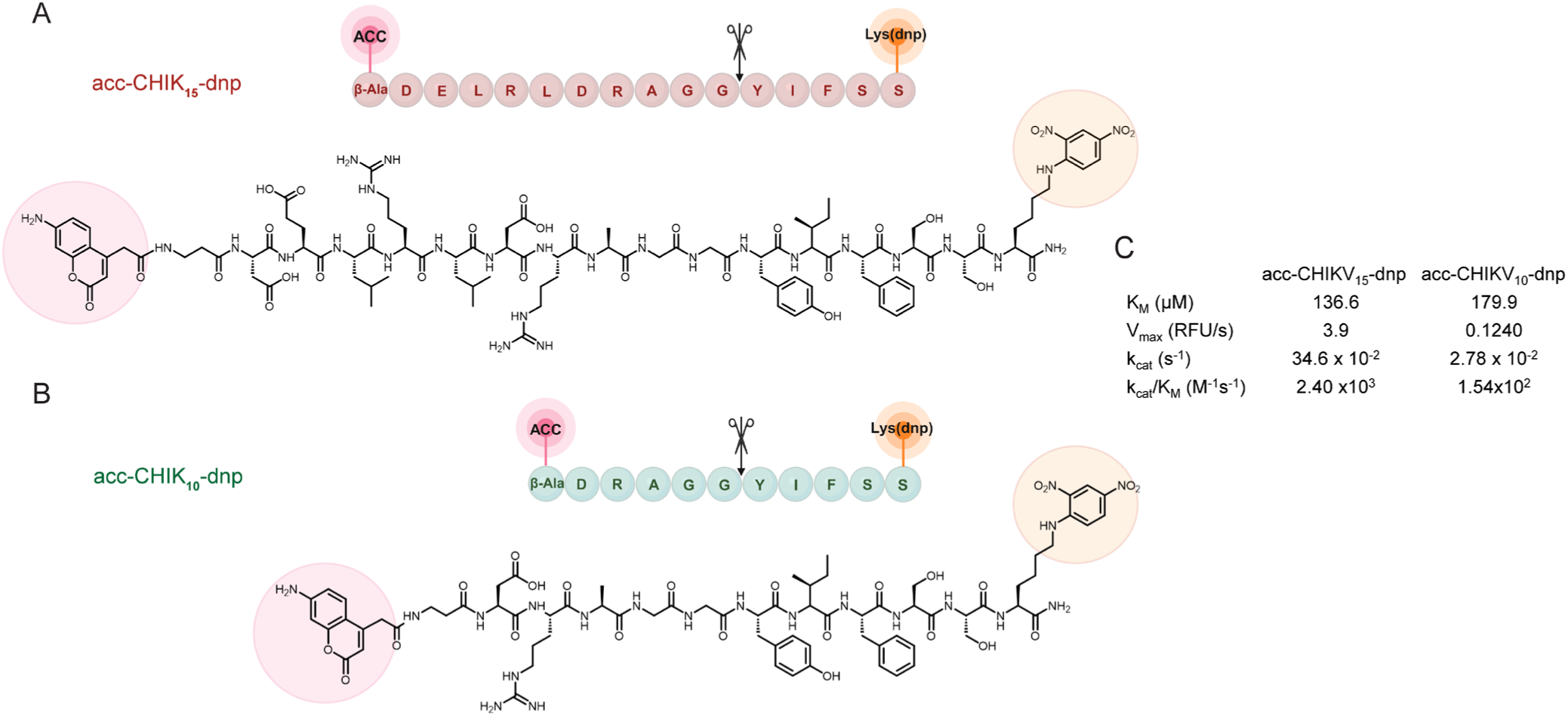
Structure of new CHIKVP substrates containing 7-amino-4-carbamoylmethylcoumarin (acc) and 2,4-dinitrophenyl-lysine (dnp) as a donor-quencher pair. **(A)** acc-CHIK_15_-dnp derives from the P10 to P5’ region of nsP3/4 cleavage site of the CHIKV polyprotein. **(B)** acc-CHIK_10_-dnp spans the P5 to P5’ region of nsP3/4 cleavage site of CHIKV polyprotein. **(C)** Kinetic parameters of CHIKVP as determined with each of these substrates. acc-CHIK_15_-dnp showed improved values than the shorter acc-CHIK_10_-dnp substrate.

### acc-CHIK_15_-dnp outperforms previously reported CHIKVP substrates

Given the catalytic parameters we observed for acc-CHIK_15_-dnp, we aimed to perform a head-to-head assessment compared to the most commonly used CHIKVP substrate, edab_8_ (Narwal et al., 2018; Saha et al., 2018; Saisawang, Saitornuang, et al., 2015; Saisawang, Sillapee, et al., 2015; Singh et al., 2018). The initial velocity (V_0_) for acc-CHIK_15_-dnp was 30 times greater than that observed for edab_8_ (Figure 4A, S6). This is even more notable given that a strong signal for acc-CHIK_15_-dnp could be observed at 10-fold lower fluorophore concentration (50 µM) than for edab_8_ which required a minimum substrate concentration of 500 µM to obtain a readily interpretable signal. We varied enzyme concentration, substrate concentration, temperature and pH to optimize the method (Figure S7). Our results are in contrast to previously reported performance of edab_8_ (Narwal et al., 2018; Saisawang, Sillapee, et al., 2015). Whereas prior reports suggested that edab_8_ generated a strong signal at concentrations as low as 1µM, in our hands, 60 µM edab_8_ was required for a readily interpretable signal (Figure S8). While this might indicate that our CHIKVP construct was less active than that used in the previous studies (Narwal et al., 2018; Saha et al., 2018; Saisawang, Sillapee, et al., 2015), it is nevertheless clear that acc-CHIK_15_-dnp produced a better signal than edab_8_ in this head-to-head comparison. acc-CHIK_15_-dnp showed more robust properties than the shorter substrates acc-CHIK_10_-dnp (both at 50 µM, Figure 4B). This suggests that CHIKVP recognition is sensitive to the length of the substrate and presence of residues on both the P and P’ sides of the cleavage site. The catalytic efficiency for acc-CHIK_15_-dnp (k_cat_/K_M_ = 2.40 ×10^3^ M^−1^s^−1^; Figure 4C) is substantially improved when compared to both our measured k_cat_/K_M_ (1.14 ×10^2^ M^−1^s^−1^) and the reported k_cat_/K_M_ (2.33 ×10^2^ M^−1^s^−1^) for edab_8_ (Narwal et al., 2018) (Figure 4C). Likewise, V_0_ at different concentrations of CHIKVP was consistently much higher for 50 µM acc-CHIK_15_-dnp than for 500 µM edab_8_ (Figure 4D, S9). This suggests that acc-CHIK_15_-dnp could be used for high throughput screening with 2.5 µM CHIKVP, which is 2-fold lower than required for edab_8_.

**Figure 4.**
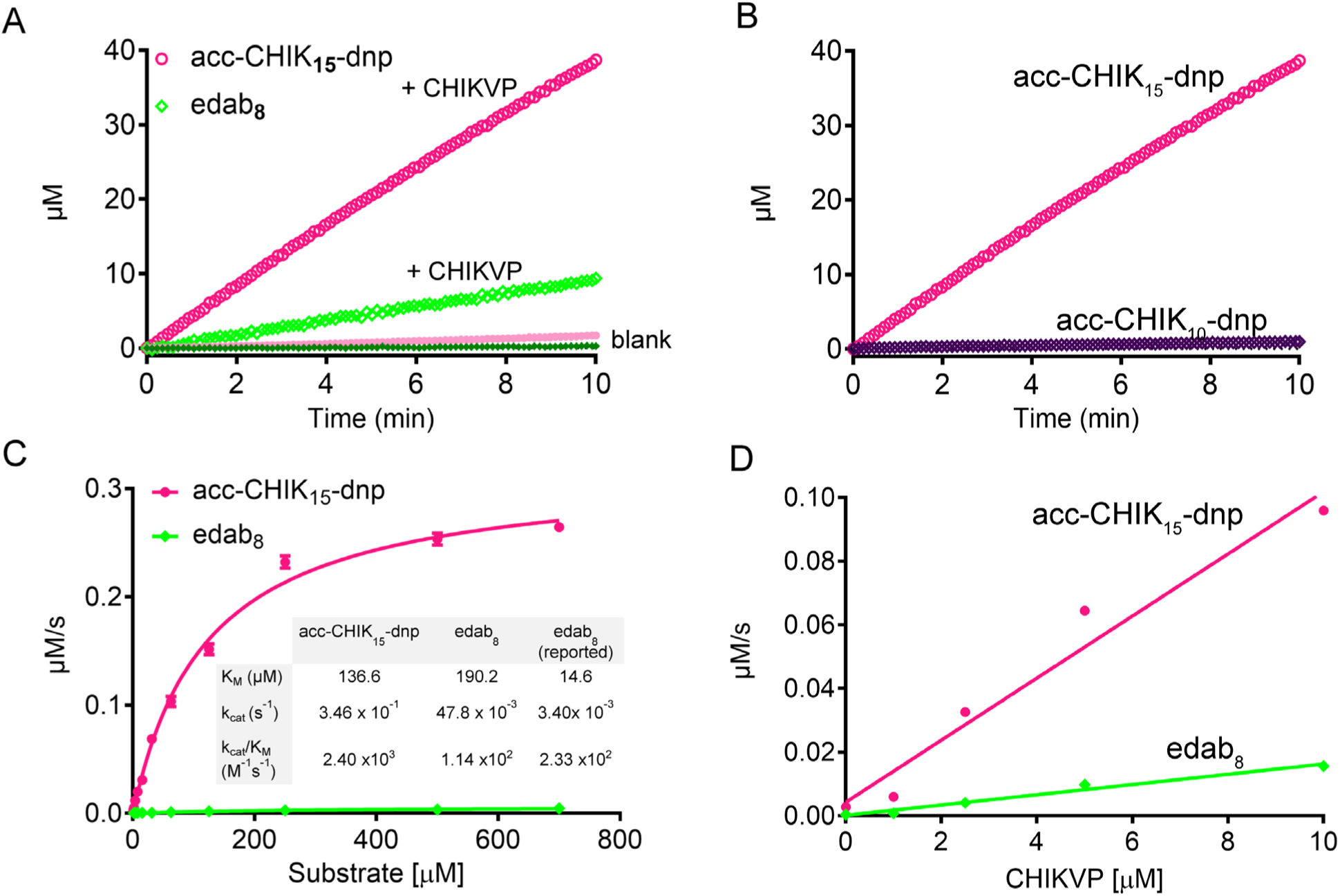
acc-CHIK_15_-dnp shows improved activity. **(A)** acc-CHIK_15_-dnp [50 µM] shows 30-fold higher V_0_ than edab_8_ [500 µM] with CHIKVP [5 µM], although 10-fold less acc-CHIK_15_-dnp is required to observe a measurable response. **(B)** The length of the peptide substrates acc-CHIK_10_-dnp and acc-CHIK_15_-dnp determines CHIKVP activity. The concentration of both substrates is 50 µM and CHIKVP is 5 µM **(C)** Substrate titration and activity of 5 µM CHIKVP monitored by acc-CHIK_15_-dnp or edab_8_. Measured kinetic parameters of CHIKVP obtained with acc-CHIK_15_-dnp or edab_8_ are compared with reported edab_8_ (Narwal et al., 2018) parameters (inset). **(D)** acc-CHIK_15_-dnp [50 µM] provides higher signal than edab_8_ [500 µM] at all CHIKVP concentrations.

### acc-CHIK_15_-dnp is optimized for CHIKVP

acc-CHIK_15_-dnp contains a peptide sequence derived from P10-P5’ from the CHIKV nsP3/4 cleavage site. The P2-P3 and P1’-P5’ residues in the nsP3/4 cleavage sites of alphaviruses CHIKV, Mayaro virus (MAYV) and Venezuelan equine encephalitis virus (VEEV) are strictly conserved (Figure 5A). In spite of the nsP3/4 sequence similarity, acc-CHIK_15_-dnp was not a robust substrate for MAYV nsP2 protease (MAYVP) or VEEV nsP2 protease (VEEVP) (Figure 5B). The catalytic site substituted variant of CHIKVP (C478A) showed no significant activity over the substrate-only control, suggesting that the presence of the protein itself does not adversely impact the assessment of proteolytic activity (Figure 5C). CHIKVP was roughly 20 times more active towards acc-CHIK_15_-dnp than the other two nsPs (Figure 5D). Although acc-CHIK_15_-dnp is optimized for CHIKVP, it is important to note that the activity of MAYVP and VEEVP can both be assessed using this substrate, albeit at a very low level. The lack of sequence identity in the P-sides of the alphaviral nsP3/4 junctions and the differential rates for cleavage of acc-CHIK_15_-dnp by CHIKVP, MAYVP and VEEVP underscore the importance of P-side residues over P’-side residues for development of effective substrates for these enzymes.

**Figure 5.**
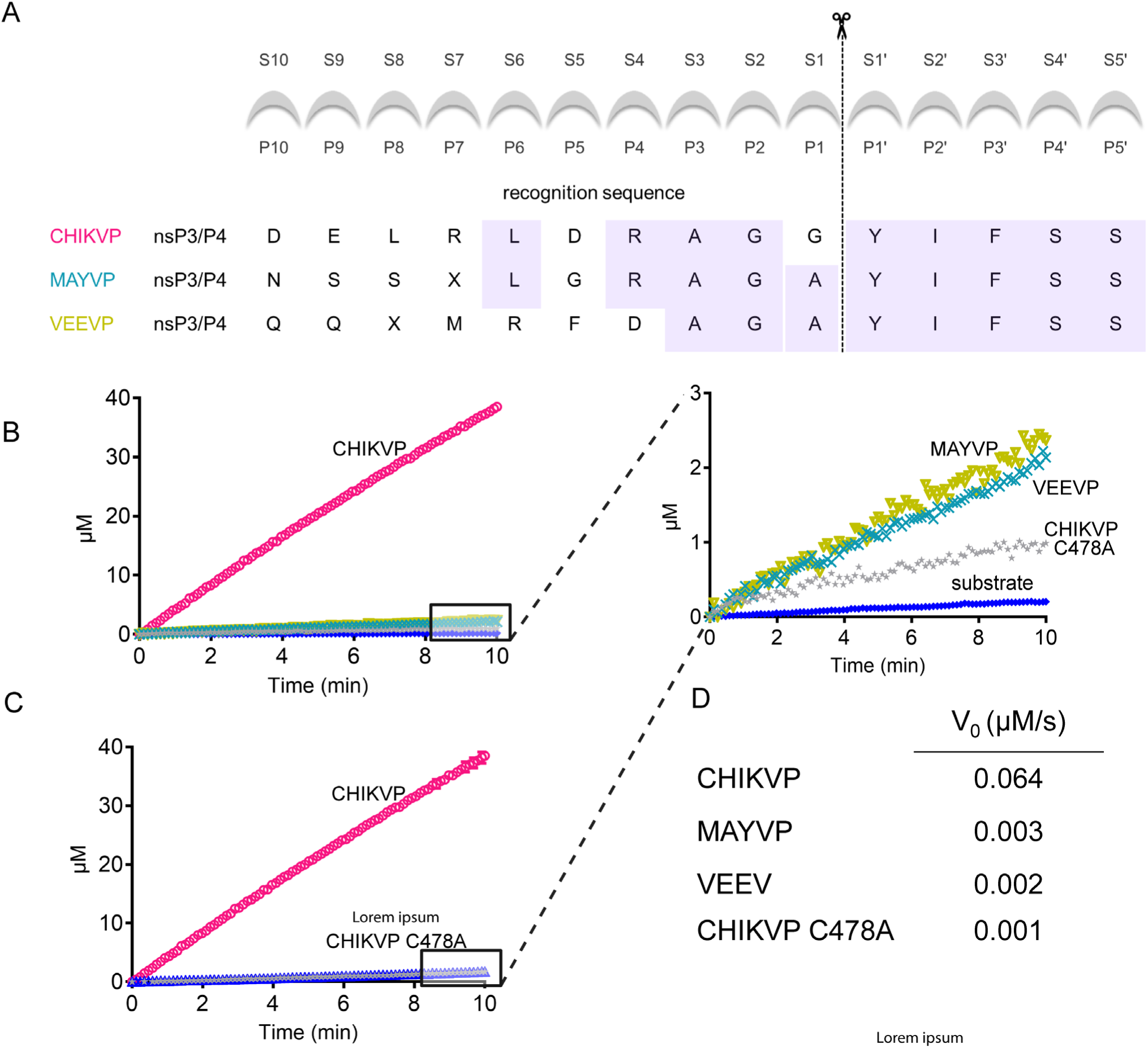
acc-CHIK_15_-dnp is specific to CHIKVP. **(A)** Aligned recognition sequences of nsP3/4 cleavage site for interrogated alphavirus proteases (MAYVP: Mayaro virus protease, VEEVP: Venezuelan equine encephalitis virus). **(B)** Comparison of activity of different alphavirus proteases (MAYVP, VEEVP) tested against acc-CHIK_15_-dnp in the fluorescence-based assay. **(C)** Inactive CHIKVP C478A in which the catalytic nucleophile has been replaced shows greatly reduced cleavage activity. **(D)** The measured initial velocities (V_0_) for alphavirus proteases from **(B, C)**.

### acc-CHIK_15_-dnp produces a robust signal for high-throughput screening

To establish effective screening protocols for small molecule inhibitors for CHIKV infection, we screened two different compound libraries using a high-throughput screening (HTS) approach using two different substrates. Initially the FDA compound library was tested against CHIKVP using the edab_8_ as the substrate. The optimized assay conditions for the HTS used 5 µM CHIKVP, 250 µM edab_8_ and 3 hours incubation at RT before assessment of inhibition (Figure 6A, Table 1). As is the industry standard for HTS, we set the hit threshold at 3σ over the mean of the library sample signals. The 3σ cutoff allowed identification of only one hit from the 960 compounds screened. This was due to the low signal-to-noise (S/N) and significant overlap (at the 2σ level) with the signal originating from edab_8_ only. The fact that the 3σ line was substantially below the background level complicates distinguishing real hits from false positives. This S/N was notably low in light of the fact that each of these compounds was assessed independently in two separate replicates, in order to obtain data that were interpretable. Thus, there was a need for a more sensitive and efficient assay for screening additional libraries to obtain reliable hits.

**Figure 6.**
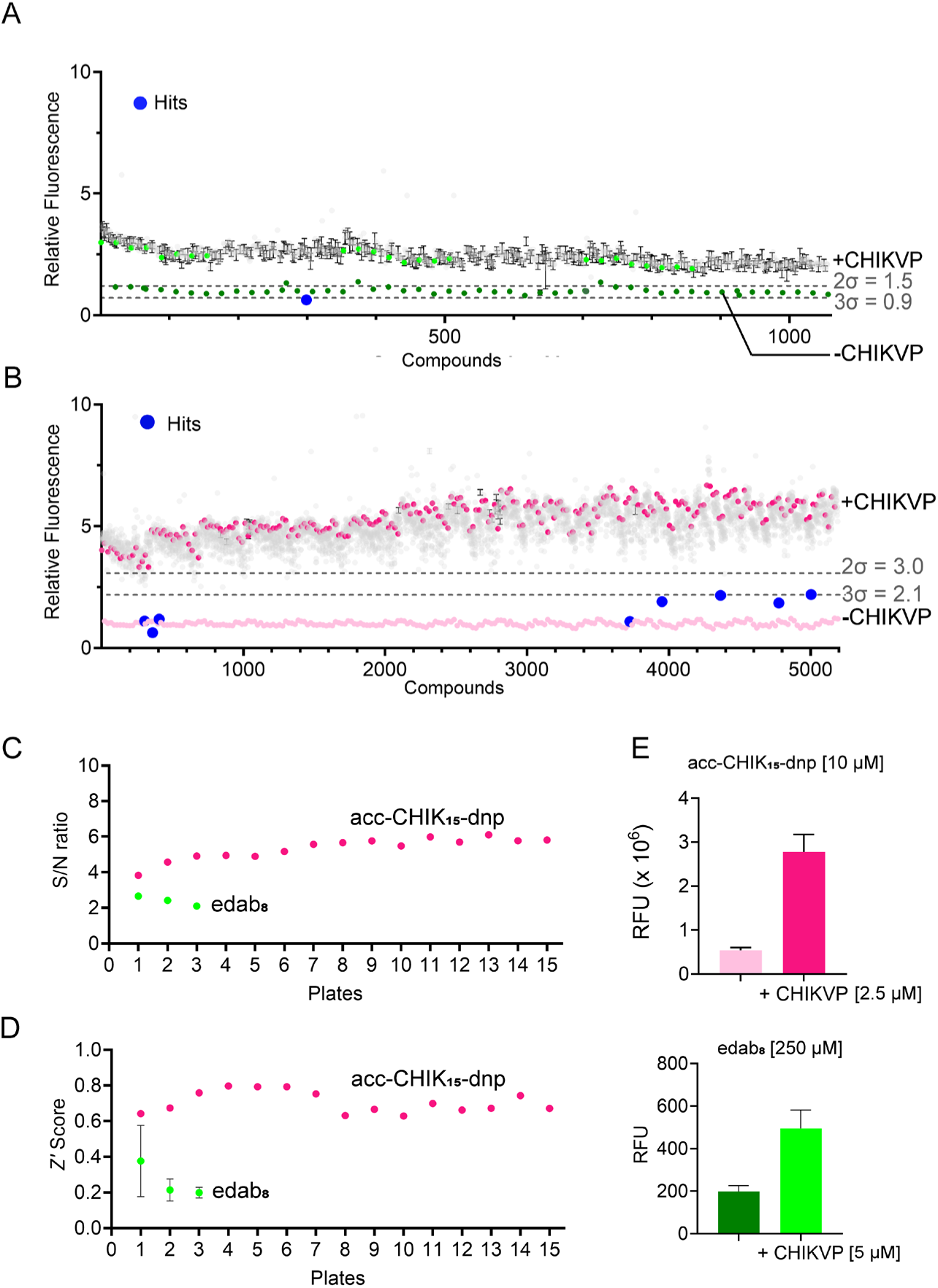
Evaluation of acc-CHIK_15_-dnp as substrate for high-throughput screening. **(A)** Screening of FDA approved compound library (25 µM) using edab_8_ as substrate (250 µM) and CHIKVP (5 µM). To obtain interpretable data, it was necessary to assay each compound in two biological replicates (n = 2). Even with replicate data, the 2*σ* cut-off is superimposable with the background signal and 3*σ* is below background indicating a poor S/N. **(B)** Screening of Life Chemicals diversity set library (50 µM) using acc-CHIK_15_-dnp as substrate (10 µM) and CHIKVP (2.5 µM) was much more readily interpretable even at lower substrate concentrations. Data from these conditions could be readily interpreted with only one biological replicate (n = 1). The hits are obtained from 3*σ* cutoff drawn above the control (substrate only) and more hits are obtained using 2*σ* values, making it a robust screening assay. **(C)** Comparison of the S/N ratios from two screening campaigns. FDA approved compound library with edab_8_ showed S/N ratios < 3 and the Life Chemicals diversity set library using acc-CHIK_15_-dnp had S/N ratios > 3. **(D)** Comparison of the *Z*’ Score ratios from two screening campaigns. **(E)** Comparison of the raw fluorescence values between the two substrates.

**Table 1.**
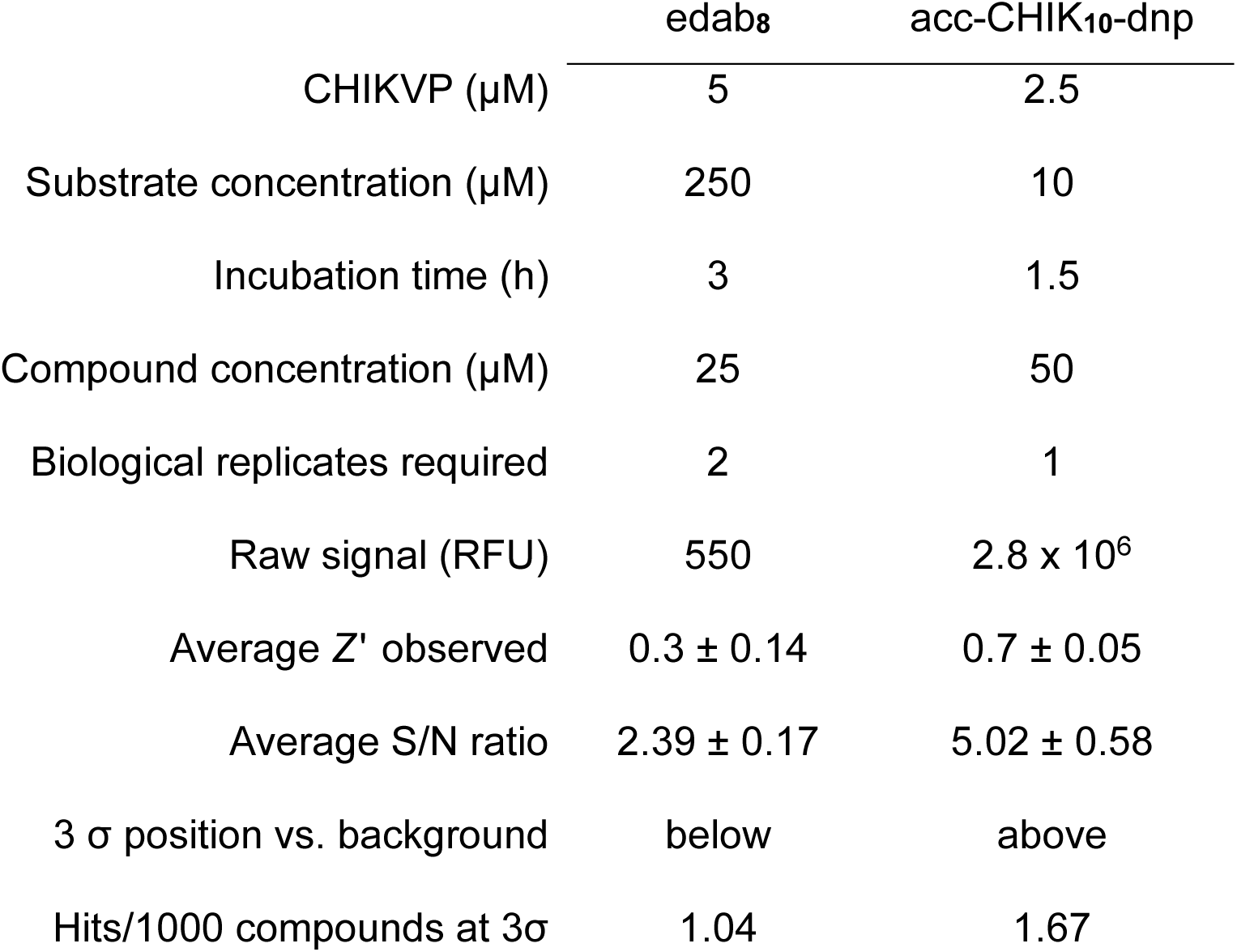
CHIKVP Screening Parameters.

We next screened LC-Diversity compound library using our newly developed acc-CHIK_15_-dnp as the substrate. Due to its improved properties, it enabled us to perform the HTS at 2.5 µM CHIKVP and 10 µM substrate with only a single replicate per compound (Figure 6B). This halving of the CHIKVP concentration provided an appreciable positive impact on the prospect of future large-scale screening. In addition, with acc-CHIK_15_-dnp the compounds were incubated only for 1.5 hours and still provided a readily interpretable fluorescence signal. From screening of 4800 compounds, eight hits were obtained using a 3σ cutoff limit. These compounds were cherry picked re-tested and found to be bona fide hits. We also performed a titration and observed IC_50_ values ranging from 1.2 to 10.6 μM (Figure S10). The high reconfirmation rate was expected since a 3σ cutoff has been reported to decrease the false-positive rate (Smith et al., 2022). In addition, the signal-to-noise ratio significantly improved for acc-CHIK_15_-dnp relative to edab_8_ (Figure 6C). The Z’ score of > 0.6 with acc-CHIK_15_-dnp (Figure 6D) further suggests that the quality of the assay was improved with the new substrate. Lastly, the fact that a substantial increase in the raw fluorescence values could be observed even at 25-fold lower acc-CHIK_15_-dnp concentration and with half the concentration of CHIKVP, underscores the improved properties of acc-CHIK_15_-dnp for HTS over edab_8_ (Figure 6E, S11). With the improved assay outcomes using acc-CHIK_15_-dnp, a peptide sequence derived from nsP3/4 cleavage site of CHIKV, we have successfully screened 4800 compounds against CHIKVP. Using our robust assay conditions, we are currently in the process of screening larger libraries to discover hits as potential CHIKVP inhibitors.

## DISCUSSION

Fluorogenic enzymatic assays are arguably the most convenient method of studying enzymatic activity. A successful assay should be able to measure enzymatic activity with high sensitivity, reproducibility and selectivity. Due to their practical applicability and rapid implementation, these assays have been widely used to screen for inhibitors of enzymatic activity on a high-throughput scale. Since each compound in HTS is typically only tested once, a good quality HTS assay relies on a good signal-to-noise ratio, a high Z’ factor (≥ 0.5) and should be able to identify the hits from false positives with a high degree of correctness (Zhang, Chung, & Oldenburg, 1999). In this study, we report a new, robust fluorogenic peptide substrate derived from nsP3/4 cleavage site of CHIKV which has been successfully used to run a HTS for the first time to discover small molecule inhibitors for CHIKVP. Our acc-CHIK_15_-dnp substrate has enabled a good quality HTS using half as much CHIKVP. Prior to this, predominantly *in-silico* screens have been reported for inhibitors of CHIKVP (Agarwal, Asthana, & Bissoyi, 2015; Das et al., 2016; Ivanova, Rausalu, Ošeka, et al., 2021; Ivanova, Rausalu, Žusinaite, et al., 2021; Kovacikova & Van Hemert, 2020; Singh et al., 2018; Souza et al., 2023) with the exception of a recent covalent fragment screen using a different substrate derived from nsP1/2 cleavage site (Merten et al., 2024). The lack of small molecule HTS attempts for CHIKVP in the past appears to have stemmed from the absence of a robust fluorogenic substrate. We anticipate that with the development of acc-CHIK_15_-dnp, additional HTS campaigns will now be undertaken, exploring a larger chemical space. This may positively contribute to the discovery of effective therapies for CHIKV, an emerging threat, with epidemic potential.

In this study, we made two significant observations which should guide the development of new alphaviral protease substrates. First, we tested the acc-CHIK_15_-dnp substrate with alphavirus proteases MAYVP and VEEP, in which the P’-side residues are strictly conserved and P4-P1 residues are conserved. In spite of this conservation, acc-CHIK_15_-dnp is only an effective substrate for CHIKVP. This clearly suggests that P-side residues are critical for efficient recognition and cleavage of substrates by alphavirus proteases. For the SFV protease (SFVP), P6-P1 residues had a major impact on cleavage (Lulla et al., 2013) Our observations likewise suggest that P6-P1 are critical for substrate recognition, but we further observe that the presence of additional residues on the P side result in more effective recognition and cleavage by CHIKVP. Second, our findings also suggest that a suitable selection of FRET donor and quencher pairs for a peptide substrate may provide a substantial enhancement of signal-to-noise ratios in performing fluorescence-based assays due to their enhanced solubility and better fluorogenic properties.

Alphaviruses are rising global public health threats (Levi & Vignuzzi, 2019) and unfortunately to date there are no antivirals available for their treatment. MAYV is one such emerging infectious disease which has symptoms similar to CHIKV (PAHO, 2019) but is yet understudied due to lack of availability of suitable substrates to probe the enzymatic activity of its nsP2 protease, which like CHIKVP, is essential to its life cycle. Interestingly, there have been no reports of small molecule inhibitor HTS campaigns using fluorogenic peptide substrates against MAYVP, VEEVP or SFV protease. The findings of this study provide a blueprint for the systematic design of fluorogenic peptide substrates for MAYVP and other alphavirus proteases in need of effective fluorogenic substrates. Such additional substrates would eventually enable HTS campaigns, supporting drug discovery efforts against these emerging viral threats.

## MATERIALS AND METHODS

Unless otherwise noted, all reagents were obtained from Fisher BioReagents^TM^.

### Expression and Purification

CHIKVP: A construct for expression of CHIKVP with an N-terminal thrombin-cleavable His_6_ tag in pET-15b plasmid was generously provided by Tom Hobman, University of Alberta. CHIKVP was transformed into BL21(DE3) electrocompetent *E. coli* and plated on ampicillin (100 µg/ mL) containing LB plates. A 50 mL *E. coli* culture was grown in LB with ampicillin (100 µg/ mL) at 37 °C overnight. 5 mL of the overnight culter was added to 1L LB containing ampicillin (100 µg/ mL) at 37 °C until the OD_600_ reached 0.6. *E. coli* were induced at this point with IPTG (Gold Biotechnology) to a final concentration of 1 mM, and maintained at 25 °C for four hours to allow protein expression. Cultures were pelleted by centrifugation and stored at −80 °C. Prior to purification the cells were thawed in lysis buffer (50 mM Tris pH 7.5, 5% glycerol, 500 mM NaCl) and lysed using a microfluidizer (Microfluidics^TM^). The lysate was clarified by centrifugation at 27,000 rcf for 50 minutes at 4 °C. The supernatant thus obtained was loaded onto a pre-equilibrated 5 mL HiTrap nickel-affinity column (GE Healthcare). After this the column was washed with wash buffer (50 mM Tris pH 7.5, 300 mM NaCl, 5% glycerol and 50 mM imidazole) before being eluted with elution buffer (50 mM Tris pH 7.5, 300 mM NaCl, 5% glycerol) using a linear gradient of 100 mM-600 mM imidazole. Protein obtained was then assessed for purity by SDS-PAGE. Fractions containing CHIKVP were pooled together. The His_6_ tag was removed by thrombin (protein: thrombin ratio 50:1) using overnight dialysis (50 mM Tris pH 7.5, 5% glycerol, 100 mM NaCl, 0.5 mM DTT (Gold Biotechnology)). The sample was filtered using a 0.22 µm syringe filter (Millipore Sigma) and reloaded onto a pre-equilibrated 5 mL HisTrap nickel-affinity column (GE Healthcare) using Buffer A (50 mM Tris pH 7.5, 5% glycerol and 0.5 mM DTT) to separate the cleaved His-tag from the CHIKVP. Although the his_6_-tag had been removed, CHIKVP bound to the 5 mL HisTrap nickel-affinity column, presumably due to ionic interactions. CHIKVP was eluted using the elution buffer (50 mM Tris pH 7.5, 300 mM NaCl, 5% glycerol) using a linear gradient of 100 mM-600 mM imidazole and the fractions were assessed for purity using SDS-PAGE (Figure S12). The pure fractions were pooled together, and their concentrations were measured by absorbance at 280 nm. The concentration of the CHIKVP was determined using Beer-Lambert’s law with extinction coefficient 50,880 M^−1^ cm^−1^ and samples were stored at −80 °C for future use.

MAYVP: A construct for expression of MAYVP with an N-terminal TEV-cleavable His_6_ -maltose binding protein (MBP) fusion tag in pET-15b plasmid was generously provided by Tom Hobman, University of Alberta. MAYVP was transformed into BL21(DE3) electrocompetent *E. coli* and plated on kanamycin (50 µg/ mL) containing LB plates. A 50 mL *E. coli* culture was grown in LB with kanamycin (50 µg/ mL) at 37 °C overnight. 5 mL of overnight culture was added to 1L LB with kanamycin (50 µg/ mL) at 37 °C until OD_600_ reached 0.6. *E. coli* were induced at this point with IPTG (Gold Biotechnology) to a final concentration of 1 mM and maintained at 25 °C for four hours to allow protein expression. Cultures were pelleted by centrifugation and stored at −80 °C. Prior to purification the cells were thawed in lysis buffer (50 mM Tris pH 8.0, 5% glycerol, 300 mM NaCl,) and lysed using a microfluidizer (Microfluidics^TM^) The lysate was clarified by centrifugation at 27,000 rcf for 50 minutes at 4 °C. The supernatant thus obtained was loaded onto a pre-equilibrated 5 mL HisTrap nickel-affinity column (GE Healthcare). After this the column was washed with wash buffer (50 mM Tris pH 8.0, 300 mM NaCl, 5% glycerol and 50 mM imidazole) before being eluted with elution buffer (50 mM Tris pH 8.0, 300 mM NaCl, 5% glycerol, using a linear gradient of 100 mM-600 mM imidazole). Protein obtained was then assessed for purity by SDS-PAGE. Fractions containing MAYVP were pooled together. The MBP tag was removed by Tabacco Etch Virus (TEV) protease (protein: TEV ratio 50:1) without shaking for 3 hours at RT. The filtered protein was then reloaded onto a pre-equilibrated 5 mL MBPTrap HP (GE Healthcare) column for further purification in Buffer A (50 mM Tris pH 8.0, 5% glycerol, 250 mM NaCl and 5 mM DTT). The protein was eluted using the elution buffer (50 mM Tris pH 8.0, 5% glycerol, 10 mM maltose using a linear gradient of 1M NaCl) and the fractions were assessed for purity using SDS-PAGE (Figure S13). The pure fractions were pooled together, and their concentrations were measured by absorbance at 280 nm. The concentration of the protein was determined using Beer-Lambert’s law with extinction coefficient 56,840 M^−1^ cm^−1^and samples were stored at −80 °C for future use.

VEEVP: A construct for expression of VEEVP with His_6_-tagged SUMO fused-VEEV nsP2 cysteine protease without the preceding helicase to form pET His6-SUMO-VEEVp_ΔC2 (His_6_-SUMO-VEEV) was constructed. VEEVP was transformed into T7 Express *lysY/I^q^* C3013 Competent *E. coli* and plated on carbenicillin-containing (100 µg/ mL) 2x Yeast Extract Tryptone agar plates. A 10 mL *E. coli* culture was grown in 2x Yeast Extract Tryptone medium with carbenicillin (100 µg/ mL) and 1% glucose at 37 °C until an OD_600_ of 0.7–0.8 was reached. 10 mL of this culture was added to 1L selection medium at 37 °C until OD_600_ reached 0.8-1.0. *E. coli* were induced at this point with IPTG (Gold Biotechnology) to a final concentration of 0.5 mM and maintained at 18 °C overnight to allow protein expression. Cultures were pelleted by centrifugation and stored at −80 °C. Prior to purification the cells were thawed in lysis buffer (50 mM Tris pH 7.0, 0.05% NP40, 0.5 mg/mL lysozyme) and sonicated (3x 1 min: pulse ON 4 sec. pulse OFF 1 sec, 50% amplitude, Branson Sonifier) followed by centrifugation at 27,000 rcf in Avanti JA-17 rotor for 45 min at 4°C. The supernatant was loaded onto a Ni-NTA resin pre-equilibrate affinity column (Qiagen (30210)) using the wash/equilibration buffer (50 mM Tris pH 7.0, 150 mM NaCl, 10 mM imidazole, and 10% glycerol), followed by washing the column with the wash buffer. Protein was eluted using an elution buffer (50 mM Tris pH 7.0, 150 mM NaCl, 300 mM imidazole, and 10% glycerol) and the fractions were assessed for purity using SDS-PAGE (Figure S14). The pure fractions were pooled together, concentrated and buffer exchanged with Corning^®^ Phosphate-Buffered Saline, 1X without calcium and magnesium, pH 7.4 ± 0.1, using Amicon^®^ ultra-4 centrifugal filter units. The purified protein was concentrated and its concentration was determined by Bradford assay. The purified protein was flash frozen for future use at −80°C.

### LC–MS Analysis Peptide Cleavage by CHIKVP

Peptide sequences DELRLDRAGG/YIFSS, DRAGG/YIFS, RAGG/YIFS (derived from nsP3/4 cleavage site of CHIKV polyprotein chain) and VEQLEDRAGA/GIIGGSRR (derived from nsP1/2 cleavage site of CHIKV polyprotein chain) were purchased from Biomatik. The substrates were dissolved in DMSO to make a 50 mM stock. For determining the cleavage of peptide substrates by CHIKVP, a total of 20 μL sample was prepared by incubating 500 μM peptide with 5 μM CHIKVP for an hour. The reaction was diluted to 7.5 µM with 0.1% formic acid to quench the reaction. Peptide samples were analyzed by liquid chromatography-mass spectrometry (LC-MS) at the UMass Amherst Mass Spectrometry Core Facility, RRID:SCR_019063. For the competition experiment, 500 µM of DELRLDRAGG/YIFSS and VEQLEDRAGA/GIIGGSRR were mixed, incubated alone and with 5 µM CHIKVP for 10 min or 30 min. 5 µL of the incubated sample was injected onto a Halo C4 2.7m, 2.1 x 150 mm column (Advanced Materials Technology) connected to an Agilent 1100 capillary liquid chromatography system with a binary pump (Agilent). The mobile phases were (A) 0.1% formic acid and (B) 0.1% formic acid in 99% acetonitrile. The system was equilibrated in 5% B at a flow rate 0.3 mL/min and the following gradient was applied: 5% B for 1 min, 5-60% B over 10 min, 60-90% B over 2 min, 90% B for 1 min. The flow was infused into a 7 T solariX FTICR mass spectrometer (Bruker) equipped with a standard electrospray source. The capillary voltage was 4.5 kV. The dry gas flow was 10 L/min, and the dry gas temperature was 250 °C. Mass spectra were acquired over 200-2,000 m/z with a 256 Kword transient size and 0.2 s accumulation time. Data were processed using DataAnalysis 5.0 (Bruker) to obtain extracted ion chromatogram (EIC) to plot the graphs. The area under the peaks was estimated using DataAnalysis 5.0 (Bruker) to calculate relative intensities of cleaved products in the competition experiment.

### NMR of CHIKVP with Native Peptide Substrates

A sample of 300 µM CHIKVP was prepared in deuterated buffer (75 mM deuterated Tris (Cambridge Isotope Laboratories, Inc) pH 7.5, 5% deuterated glycerol (Cambridge Isotope Laboratories, Inc) and 10% D_2_O (Cambridge Isotope Laboratories, Inc)). Peptide solutions in deuterated DMSO (Cambridge Isotope Laboratories, Inc) were added to a final concentration of 1.5 mM and incubated for 10 minutes. Aliquots of 170 μL were transferred into standard 3mm NMR tubes for NMR measurements. All NMR spectra were acquired on a Bruker AVANCE III solution-state 600 MHz NMR spectrometer equipped with a liquid helium-cooled QCI (H/F, C, N, P) cryoprobe. All NMR data were collected at T=298K. All pulse programs used for data collection were from Topspin (version 3.6.2, Bruker). 1D-^1^H experiments were collected with excitation sculpting solvent suppression (Hwang & Shaka, 1995). All data were collected and processed using Topspin.

### Synthesis of Fluorogenic Substrates

Synthesis of the substrates with ACC-Lys (dnp) donor:quencher pair was carried out using a Liberty Blue Automated Peptide Synthesizer, following standard 9-fluorenylmethoxycarbonyl (Fmoc) solid-phase peptide chemistry at a 0.1 mmol scale. A four-fold excess of amino acids was used relative to the pre-loaded Rink Amide (RA) resin (particle size: 100–200 mesh, loading: 0.7 mmol/g) (Iris Biotech GmbH). Each amino acid (Combi-Blocks) was coupled in N,N-dimethylformamide (DMF) using diisopropylcarbodiimide (DIC) (1 equiv.) (Iris Biotech GmbH Marktredwitz, Germany) and Ethyl cyano(hydroxyimino)acetate (Oxyma) (2 equiv.) (Merck) as coupling reagents. The Fmoc protecting group was removed with 20% piperidine (Iris Biotech GmbH) in DMF (Avantor). After the aspartic acid coupling, the resin was transferred to a glass reaction vessel, and Fmoc-ACC-βAla-OH (2.5 eq) (Combi-Blocks) was coupled in DMF using 2-(1H-7-azabenzotriazol-1-yl)-1,1,3,3-tetramethyluranium hexafluorophosphate (HATU) (2.5 equiv.) (Iris Biotech GmbH) and 2,4,6-trimethylpyridine (2.5 equiv.) (Merck) as coupling reagents. The reaction mixture was gently shaken for 24 hours at room temperature. After this time, the resin was washed three times with DMF, and the reaction was repeated using 1.5 eq of the same reagents to improve the yield of Fmoc-ACC-βAla-OH coupling to the resin. After another 24 hours, the resin was washed with DMF and the Fmoc protecting group was removed using 20% piperidine in DMF (5, and 25 min). Subsequently, the resin was washed with dichloromethane (DCM) (Avantor) three times and methanol (MeOH) (Avantor) three times, then dried over P_2_O_5_ (Avantor). After the synthesis was completed, the peptides were cleaved from the resin with a mixture of cold TFA (Iris Biotech GmbH): TIPS (Merck): H_2_O (%, v/v/v 95:2.5:2.5; 2 hours, shaking once per 15 min). The substrates were precipitated in cold diethyl ether, centrifuged, and purified by HPLC. The purity of each substrate was confirmed with an analytical UHPLC system using a Sepax Bio-C18 3 µm 200 Å column (150 x 2.1 mm). The solvent composition was as follows: phase A (water/0.1% formic acid) and phase B (acetonitrile/0.1% formic acid); gradient, from 5% B to 95% B over a period of 15 min, flow rate 0.3 ml/min. The molecular weight of each substrate was confirmed by mass spectrometry using an LCMS-2050 SQ Shimadzu with Heated Dual Ion Source (Heated DUIS™) and single quadrupole module. Mass characterization peaks were obtained at 819.83 Da [M+2H]^2+^, and 1638.92 Da [M+H]^+^, for acc-CHIK_10_-dnp and 1133.14 Da, [M+2H]^2+^ for acc-CHIK_15_-dnp.

Synthesis of the substrate with Abz-Y(NO_2_) donor quencher pair was carried out using a Liberty Blue Automated Peptide Synthesizer, following standard 9-fluorenylmethoxycarbonyl (Fmoc) solid-phase peptide chemistry at a 0.1 mmol scale. A two-fold excess of amino acids was used relative to the pre-loaded Rink Amide (RA) resin (particle size: 100–200 mesh, loading: 0.8 mmol/g) (Novabiochem^®^). Each amino acid (Millipore Sigma) was coupled in N,N-dimethylformamide (DMF) using diisopropylcarbodiimide (DIC) (1 equiv.) (Millipore Sigma) and Ethyl cyano(hydroxyimino)acetate (Oxyma) (2 equiv.) (Millipore Sigma) as coupling reagents. The first amino acid serine was coupled manually using the same steps described above. The Fmoc protecting group was removed with 20% piperidine in DMF (Fisher chemicals^TM^). After the aspartic acid coupling, Boc-2-amino benzoic acid (ABz) (3.0 eq) (Chem Impex) was coupled in DMF using 2-(1H-7-azabenzotriazol-1-yl)-1,1,3,3-tetramethyluranium hexafluorophosphate (HATU) (Millipore Sigma) and 2,4,6-trimethylpyridine (Millipore Sigma) as coupling reagents. Subsequently, the resin was washed with dichloromethane (DCM). After the synthesis was completed, the peptides were cleaved from the resin with a mixture of cold TFA (Millipore Sigma): TIPS (Millipore Sigma) : H_2_O (%, v/v/v 95:2.5:2.5; 3 hours). This step also aided the removal of Boc protection group. The substrate was precipitated in cold diethyl ether, centrifuged, purified by HPLC and characterized using mass spectrometry on MALDI-TOF/TOF UltrafleXtreme (Bruker) with the mass characterization peak at 1197.53 Da.

### Determination of Activity of CHIKVP and VEEVP

The FRET assays to determine the activity of CHIKVP were carried out in 384-well plates (Corning 3575) in 20 µL reaction volumes. For the acc-CHIK_15_-dnp substrate the final concentration was 50 µM and CHIKVP enzyme was diluted in 50 mM Tris pH 7.5, 1mM CHAPS, to the desired concentration. To ensure that no soluble aggregates of acc-CHIK_15_-dnp were present, we measured the solubility of the substrate (500 µM) in activity assay buffer by ^1^H-NMR with TSP as internal standard. We also observed no aggregates by eye (Figure S15). The activity was measured in duplicate at 25 °C. The fluorescence was read (λ_ex_ = 340 nm and λ_em_ = 490 nm) using a SpectraMax M5 spectrometer. For all other substrates the final substrate concentration was 500 µM, with CHIKVP at 5 µM in a 20 µL reaction volume. Each measurement was performed in duplicate in a set of three independent experiments. The activity assay was optimized using varied enzyme concentration, substrate concentration, temperature and pH. Reaction progress curves and other titrations were plotted using GraphPad Prism Software. For enzyme kinetics, similar reaction volumes were used with varying concentrations of the substrate. To directly compare V_0_, 1mM of acc-CHIK_15_-dnp and edab_8_ (Biomatik) were fully digested using trypsin (substrate: trypsin ratio 100:1) followed by serially diluting these fully cleaved substrates to obtain standard curves (Figure S6).The solubility of acc-CHIK_15_-dnp substrate was assessed using DLS measurements at 10 different concentrations in the 50 mM Tris pH 7.5, 1mM CHAPS buffer.

The FRET assay to determine the activity of VEEVP was carried out in 384-well plates (Corning 3575) in 20 µL reaction volumes. For (TF5-C-LQEAGA/GSVE-K-(TQ5)-M) substrate the final concentration was 25 µM and VEEVP enzyme was diluted in 1X PBS, 0.01% Triton X, 5 mM DTT to the final concentration of 5 µM. The activity was measured in duplicate at 25 °C in a set of three independent experiments. The fluorescence was read (λ_ex_ = 640nm and λ_em_ = 670 nm) using a SpectraMax M5 spectrometer for 10 min (Figure S16). Reaction progress curves were plotted using GraphPad Prism Software.

### High-Throughput Screen Assay Development

Screening of FDA Compound Library with CHIKVP. The FDA-approved library purchased from Microsource US and International Drug Collection consisted of 1280 compounds of which 960 compounds were screened at a final conc. of 25 µM in 10 µL final volumes in duplicate. Each plate was read twice. Briefly, 50 nL of compound stock (from a 5 mM compound stock in DMSO) was transferred into wells of a 384-well plate (Corning 4514) using an Echo 650 (Beckman Coulter). 50 nL DMSO was added to control wells (substrate only and substrate with CHIKVP). CHIKVP (44 µM stock) was diluted to 10 µM in buffer (50 mM Tris pH 7.5, 1 mM CHAPS, 10 mM DTT), and 5 µL was dispensed into each well to a final CHIKVP concentration of 5 µM, excluding the substrate control wells, to which, 5 µL of buffer was added. Plates were incubated at room temperature for 1 hour to allow compound binding to CHIKVP, after which, 5 µL of edab_8_ (Biomatik) (0.5 mM diluted in buffer) was added to all the wells (final conc. 0.25 mM). All the plates were then incubated at room temperature for 3 hours. The evolved fluorescence was measured (λ_ex_= 320 nm, λ_em_= 490 nm) using an EnVision 2105 (PerkinElmer). The data were normalized to the substrate only samples in each plate. S/N ratios were calculated by dividing the average signal from wells containing both substrate and CHIKVP to the average signal from the substrate only wells. The Z’ factor was calculated using the following formula:

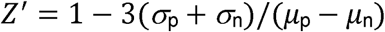

where, µ is the mean and σ is the standard deviation for both positive (p) and negative (n) controls. The positive control (p) in this experiment was CHIKVP with substrate and the negative control (n) was substrate only.

Screening of LC Diversity Set compound library with CHIKVP. The LC Diversity Set purchased from Life Chemicals consisted of 5120 compounds of which 4800 compounds were screened at a final concentration of 50 µM in a 10 µL final volume in a single replicate in a manner analogous to the FDA compound library, with the following changes: CHIKVP (36.2 µM stock) was diluted to 2.8 µM in buffer (50 mM Tris pH 7.5, 1 mM CHAPS), and 9 µL was dispensed to each well (final conc. 2.5 µM), excluding the substrate control wells, to which, 9 µL of buffer was added. Plates were incubated at room temperature for 1 hour, after which, 1 µL of acc-CHIK_15_-dnp substrate (GenScript) (0.1 mM diluted in buffer) was added to all the wells (final conc. 0.01 mM). All the plates were incubated at room temperature for 1.5 hours, and fluorescence was measured (λ_ex_ = 340 nm, λ_em_ = 480 nm). The hits thus obtained were tested for inhibition by two-fold serial dilution of the compound (starting at 50 µM) with CHIKVP (5 µM). After 1 hour incubation inhibition was checked using fluorescence activity assay. The IC_50_ curves were plotted using GraphPad Prism Software.

## Supporting information

Supplemental figures for A Robust Flurogenic Substrate for CHIKVP nsP2

## ABBREVIATIONS

ACC: 7-amino-4-carbamoylmethylcoumarin
CHIKV: Chikungunya virus
CHIKVP: Chikungunya virus protease
HTS: high throughput screening
Lys(dnp): 2,4-dinitrophenyl-lysine
MAYV: Mayaro virus
MAYVP: Mayaro virus nsP2 protease
nsP2: non-structural protein 2
SFV: Semliki Forest Virus
SFVP: Semliki Forest Virus Protease
S/N: signal-to-noise
VEEV: Venezuelan equine encephalitis virus
VEEVP: Venezuelan equine encephalitis virus nsP2 protease

## SUPPORTING INFORMATION

- Activity of CHIKVP to cleave Boc-RGG-amc, Ac-LRGG-acc, ISG15-amc, ISG15-Rh110, and ABz-RAGGYIFY(NO_2_)S).
- Mass spectral analysis of cleavage of peptide substrates DELRLDRAGG/YIFSS, DRAGGYIFS and VEQLEDRAGA/GIIGGSRR studied with CHIKVP using LC-MS.
- NMR timescale to peak shift relationship
- Activity of CHIKVP at varying enzyme and acc-CHIK_15_-dnp or edab_8_ concentrations.
- IC_50_ curves for hits obtained from high throughput screening.
- Solubility assessment of acc-CHIK_15_-dnp.
- Purification SDS-PAGE gels for CHIKVP, MAYVP and VEEVP.
- Activity of VEEVP.

## AUTHOR CONTRIBUTIONS

**Sparsh Makhaik:** Developed and executed CHIKVP nsp2 purification protocols, CHIKVP and MAYVP protein purification, experimental setup, assay development, designing substrates and synthesizing substrate with ABz-Y (NO_2_) pair, writing original draft, data curation, review and editing. **Wioletta Rut:** Design and synthesis of peptide substrate with ACC-Lys (dnp) FRET pair. **Shruti Choudhary:** HTS assay development, experimental setup, data analysis and HTS data curation. **Tulsi Upadhyay:** VEEVP protease purification, experimental setup and assay development. **Chenzhou Hao:** designing and characterizing the VEEVP substrate. **Michael Westberg:** creating the VEEVP nsp2 bacterial expression plasmid and protocols for protease purification. **Cedric Bobst:** Performing the LC-MS experiments. **Euna Yoo:** providing ISG15 substrates for activity assays. **Jasna Fejzo:** Providing NMR pulse sequences, running NMR experiments and interpreting NMR results. **Michael Z Lin:** advising on VEEVP substrate design. **Paul Thompson:** advising on HTS assay and interpretation of HTS results. **Matthew Bogyo:** Advising on VEEVP assays. **Marcin Drag:** advising on CHIKVP susbtrate design with ACC-Lys (dnp) FRET pair. **Jeanne A. Hardy:** Conceptualization; funding acquisition; overall supervision; project administration; writing original draft; review and editing; data curation.

## ACKNOWLEDGEMENTS

We thank Tom Hobman, University of Alberta for providing us with the plasmids for Chikungunya virus protease and Mayaro virus protease. Sparsh Makhaik was funded by a Translational Graduate Student Assistantship from the Institute for Applied Life Sciences, UMass Amherst. The Hardy lab was supported by Canadian Institute of Health Research PS 162417. The Drag laboratory is supported by the “TEAM/2017-4/32” project, which is conducted within the TEAM program of the Foundation for Polish Science cofinanced by the European Union under the European Regional Development Fund. Figures in this manuscript were Created in BioRender.com.

