## Supplemental figures for A Robust Flurogenic Substrate for CHIKVP nsP2 for "A Robust Fluorogenic Substrate for Chikungunya Virus Protease (nsP2) Activity"

Contents


| | | Page no. |
|---|---|---|
| 1. | Activity of CHIKVP with Boc-AGG-amc and Ac-LRGG-amc substrates | S1 |
| 2. | Activity of CHIKVP with ISG15 and ABz-RAGGYIFY($NO_2$)S substrates | S2 |
| 3. | Mass Spectra for cleavage of DELRLDRAGG/YIFSS, DRAGGYI and VEQLEDRAGA/GIIGGSRR | S3-S4 |
| 4. | LC-MS quantitation for DELRLDRAGG/YIFSS and VEQLEDRAGA/GIIGGSRR with CHIKVP | S4 |
| 5. | Relationship of NMR peak shift to exchange regime | S5 |
| 6. | Standard curve of substrates with trypsin digest | S5 |
| 7. | Activity of CHIKVP with acc-CHIK$_{15}$-dnp at different pH and temperature | S6 |
| 8. | Activity of CHIKVP with varying concentrations of edab$_8$ | S6 |
| 9. | Activity curves for varying enzyme and acc-CHIK$_{15}$-dnp concentration for CHIKVP | S7 |
| 10. | $IC_{50}$ curves for hits obtained from high throughput screening | S7 |
| 11. | $V_0$ graphs for acc-CHIK$_{15}$-dnp and edab$_8$ with different CHIKVP concentration over different substrate concentrations | S8 |
| 12. | Purification gels for CHIKVP | S8-S9 |
| 13. | Purification gels for MAYVP | S9-S10 |
| 14. | Purification gels for VEEVP | S10 |
| 15. | Solubility assessment of acc-CHIK$_{15}$-dnp | S11 |
| 16. | Activity of VEEV | S11 |

A

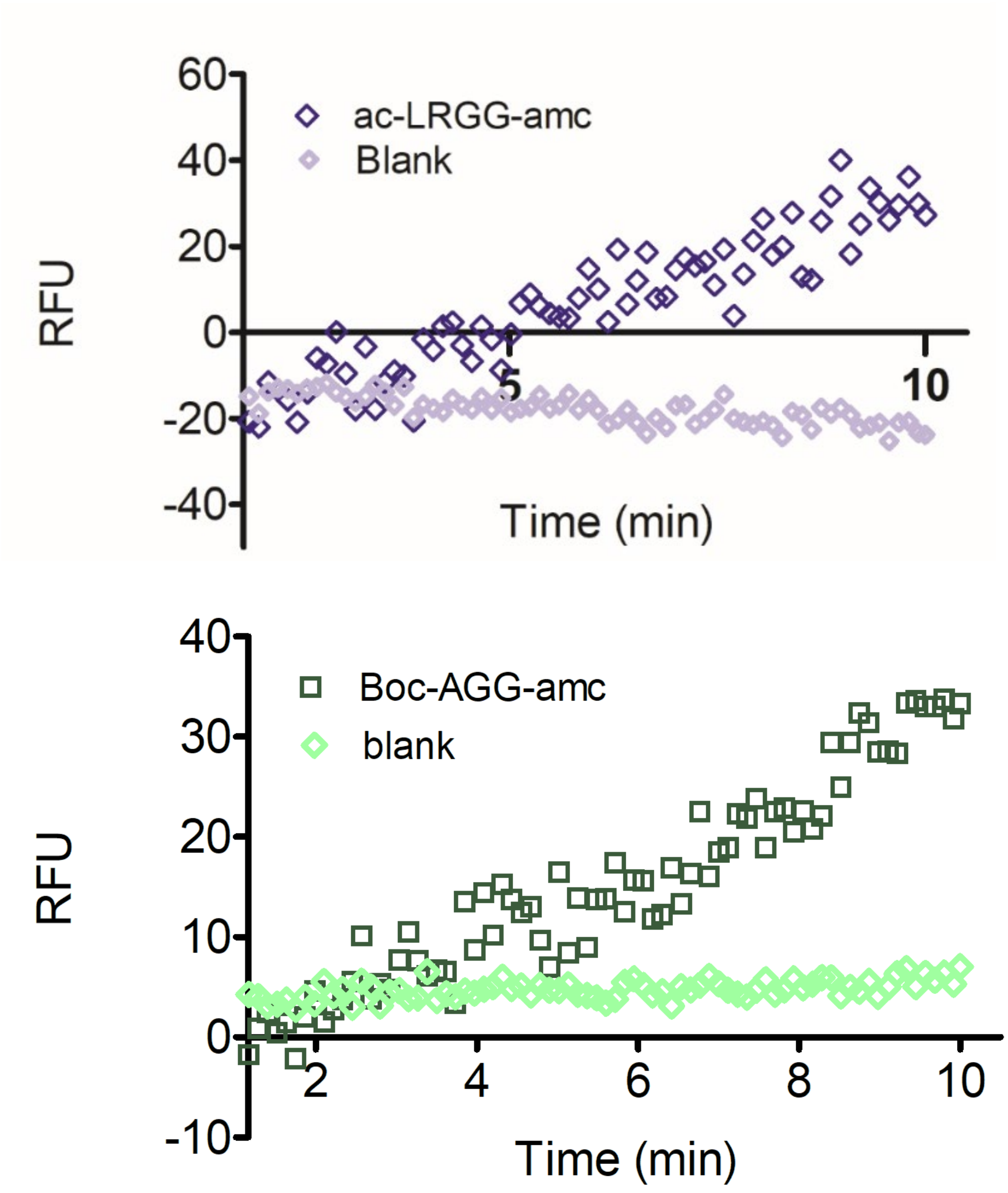


**Figure S1.** Fluorescence based activity assay for CHIKVP (A) with ac-LRGG-amc (B) Boc-AGG-amc where amc is 7-amino-4-methylcoumarin.

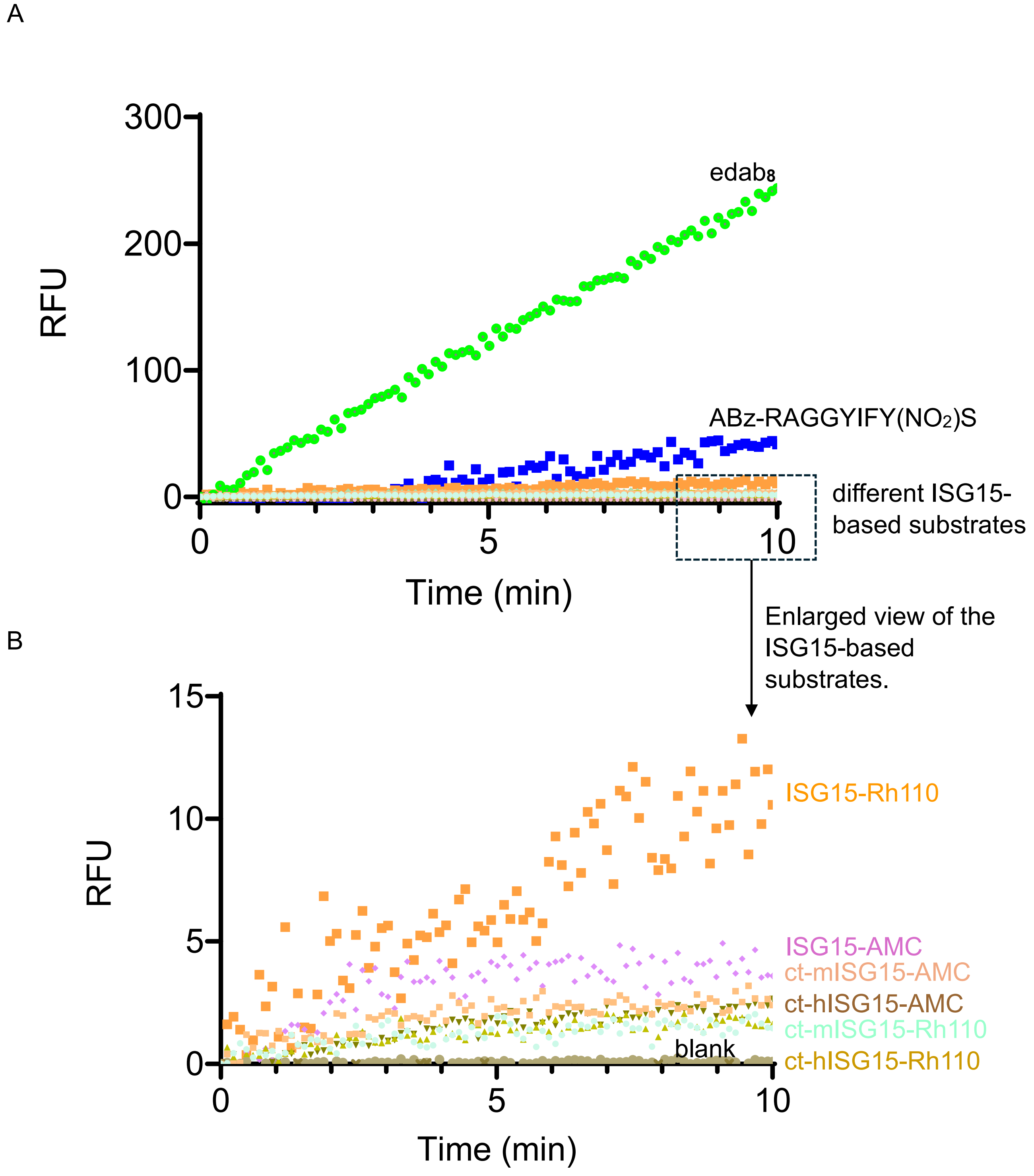


**Figure S2.** (A) Activity curves of different ISG15-based fluorogenic substrates and ABz-RAGGYIFY($NO_2$)S tested against CHIKVP and compared with edab$_8$ (B) Enlarged view of different ISG15-based substrates when tested against CHIKVP. None of these substrates showed enhanced activity when compared to edab$_8$, the most widely used substrate in literature.

A

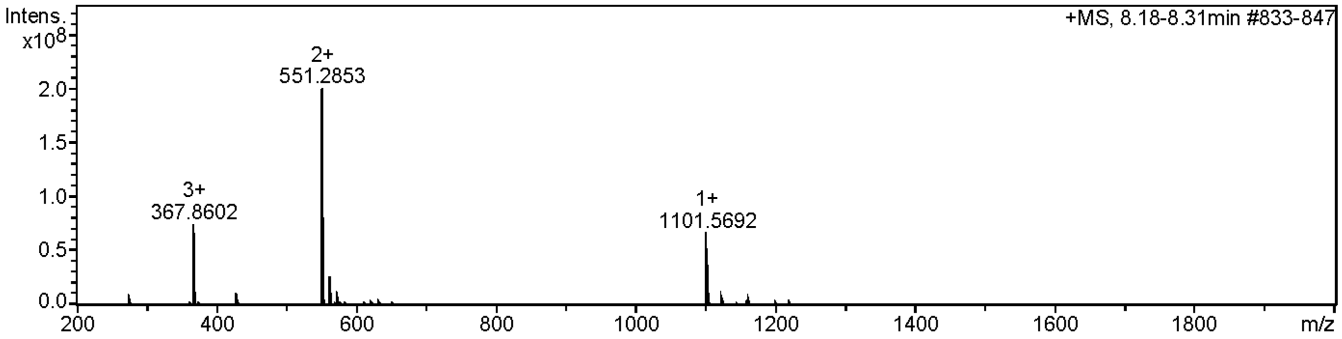
Intens.
x10^8
+MS, 8.18-8.31min #833-847
3+
367.8602
2+
551.2853
1+
1101.5692
2.0
1.5
1.0
0.5
0.0
200
400
600
800
1000
1200
1400
1600
1800
m/z

B

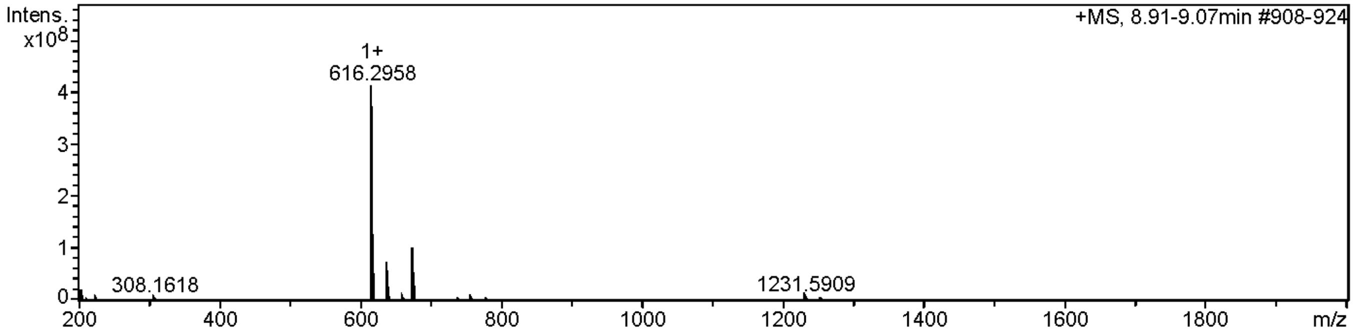
Intens.
x10^8
+MS, 8.91-9.07min #908-924
1+
616.2958
308.1618
1231.5909
4
3
2
1
0
200
400
600
800
1000
1200
1400
1600
1800
m/z

C

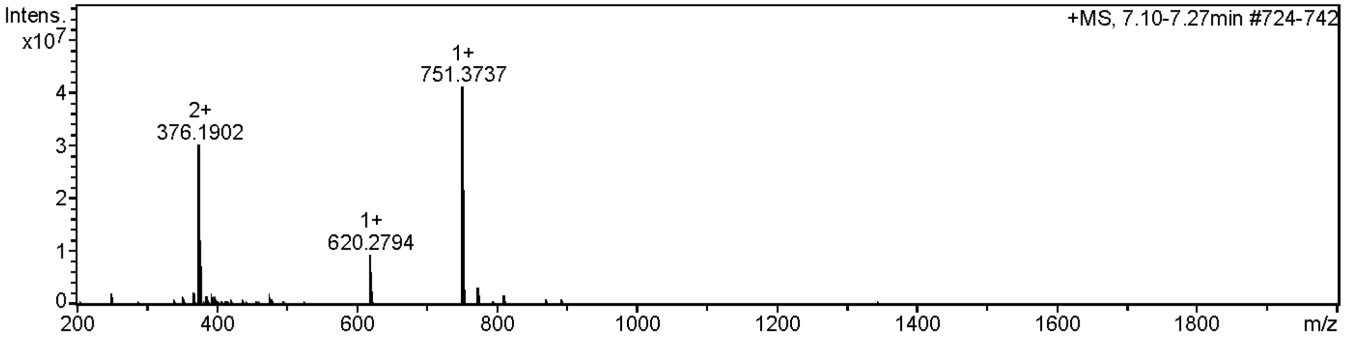
Intens.
x10^7
+MS, 7.10-7.27min #724-742
2+
376.1902
1+
620.2794
1+
751.3737
4
3
2
1
0
200
400
600
800
1000
1200
1400
1600
1800
m/z

D

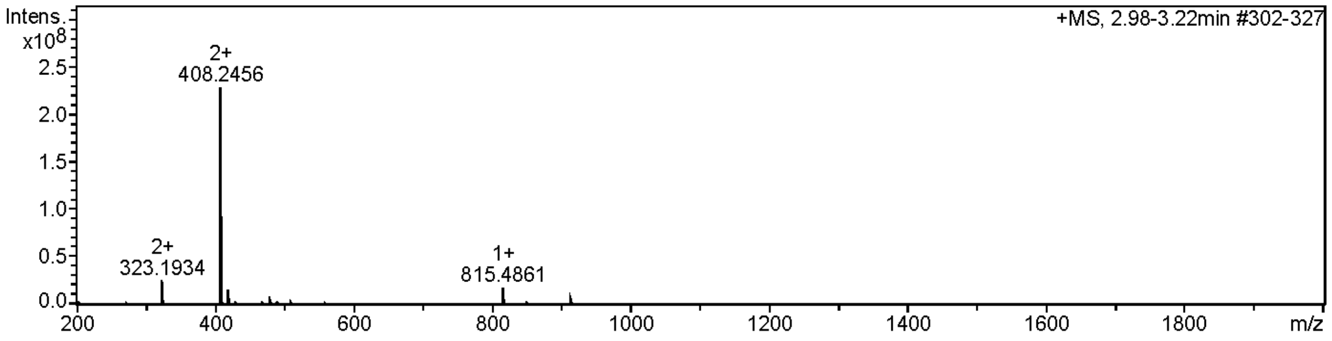
Intens.
x10^8
+MS, 2.98-3.22min #302-327
2+
323.1934
2+
408.2456
1+
815.4861
2.5
2.0
1.5
1.0
0.5
0.0
200
400
600
800
1000
1200
1400
1600
1800
m/z

E

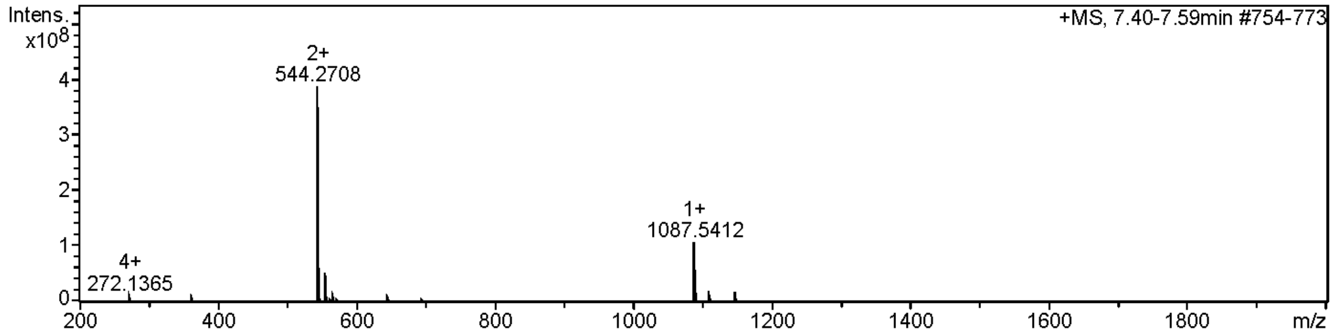


**Figure S3.** (A, B) Mass spectra for cleavage of DELRLDRAGG/YIFSS where (A) Mass spectrum for fragment DELRLDRAGG (B) Mass spectrum for fragment YIFSS (C) Mass spectrum for cleavage assay on DRAGG/YI. No cleavage of this peptide was observed (D, E) Mass spectra for cleavage for VEQLEDRAGA/GIIGGSRR where (D) Mass spectrum for fragment GIIGGSRR (E) Mass spectrum for fragment VEQLEDRAGA.

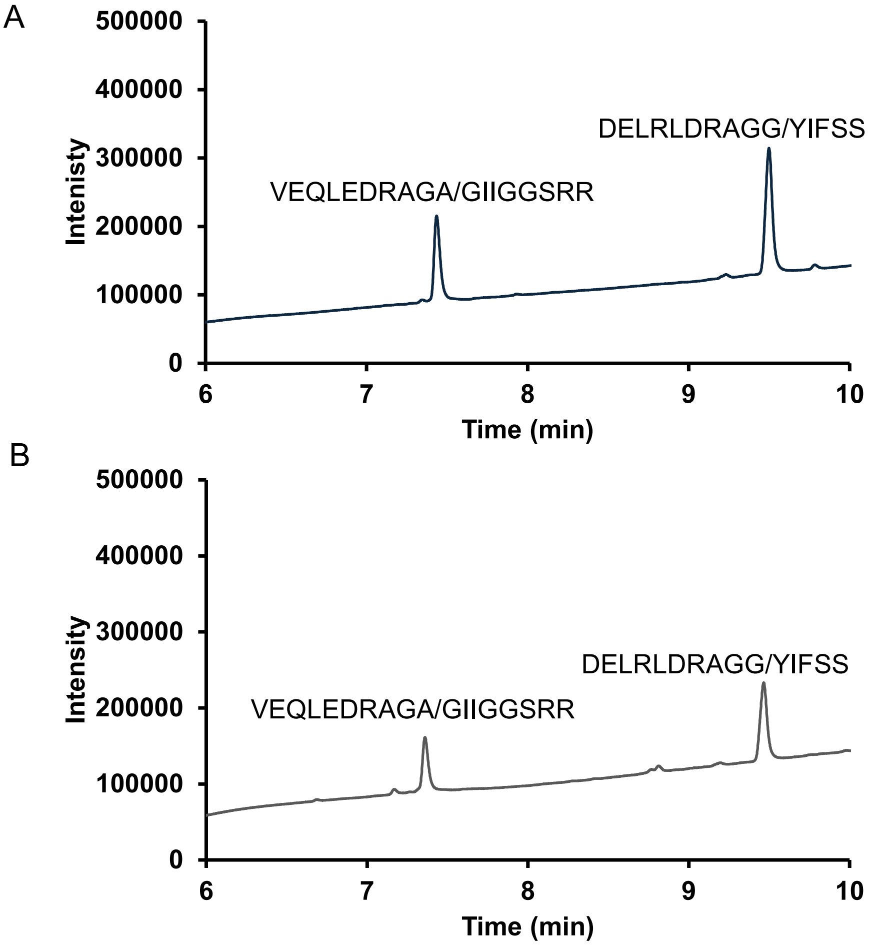


**Figure S4.** UV graphs for LC-MS quantitation of competition assay between DELRLDRAGG/YIFSS and VEQLEDRAGA/GIIGGSRR peptides with CHIKVP. (A) At 0 min without CHIKVP (B) After 10 min with CHIKVP.

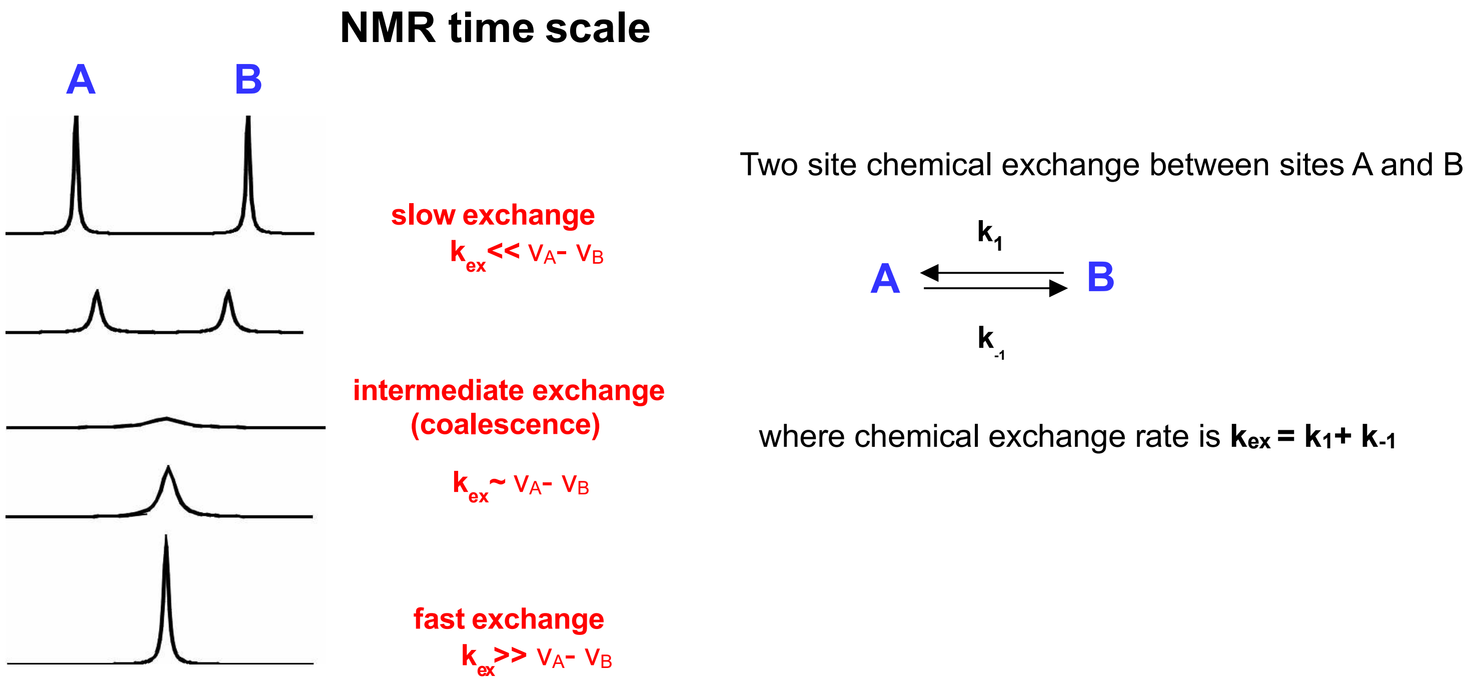


**Figure S5.** Two site chemical exchange between A and B. Site A peak resonates at $v_A$ and site B at $v_B$ frequency in the NMR spectrum. Chemical exchange in the NMR time scale can be slow when exchange rate $k_{ex} << v_A - v_B$, intermediate when $k_{ex} \sim v_A - v_B$ or fast when kex >> $v_A - v_B$.

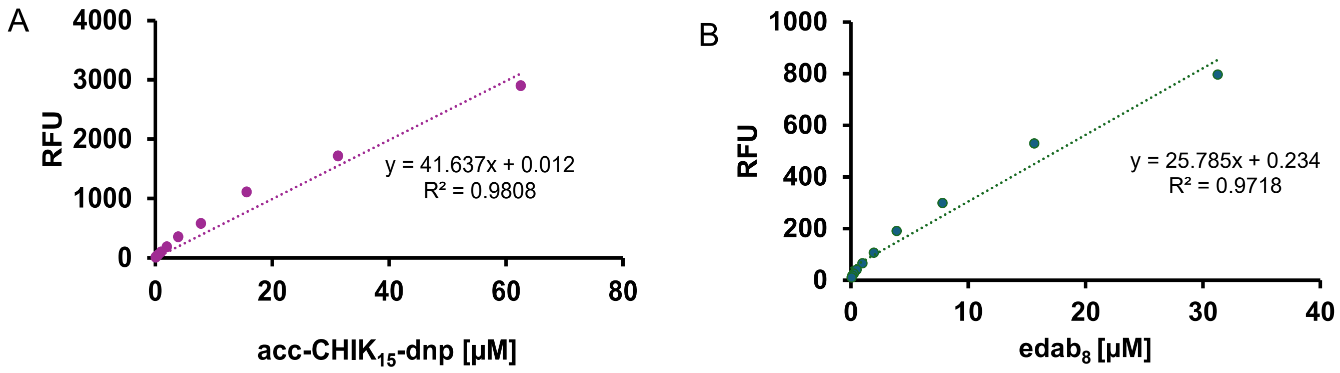


**Figure S6.** Standard curve of trypsin digest of (A) acc-CHIK$_{15}$-dnp substrate (B) edab$_8$ substrate.

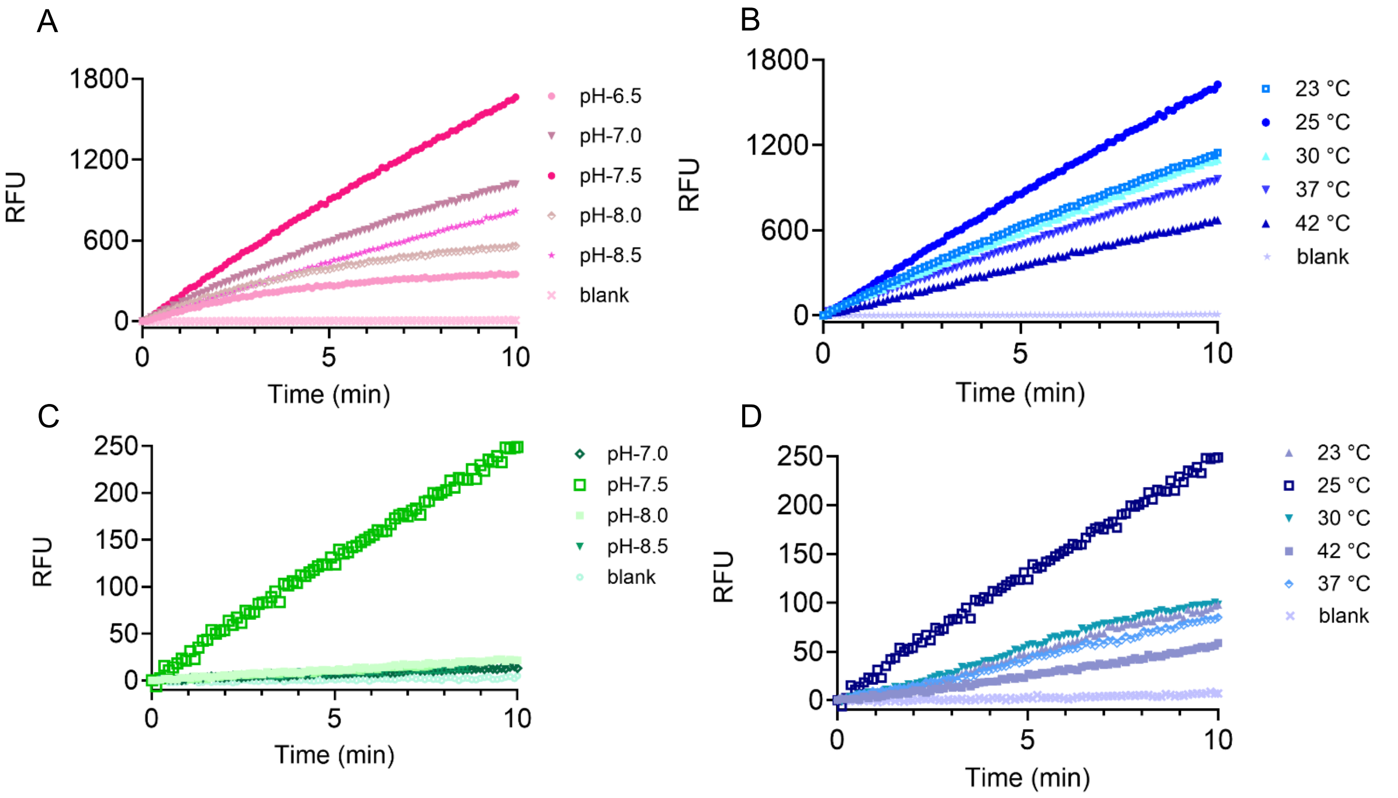


**Figure S7.** Activity 5 µM of CHIKVP as observed at different pH and temperatures. (A, B) with 50 µM acc-CHIK$_{15}$-dnp substrate (C, D) with 500 µM edab$_8$ substrate.

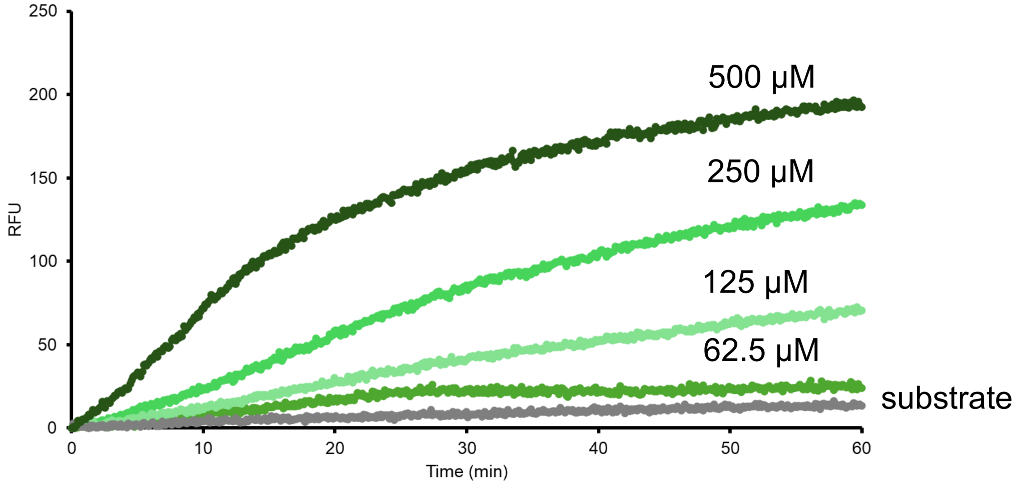


**Figure S8.** Fluorescence based activity assay for 5 µM CHIKVP with edab$_8$ at varying concentrations.

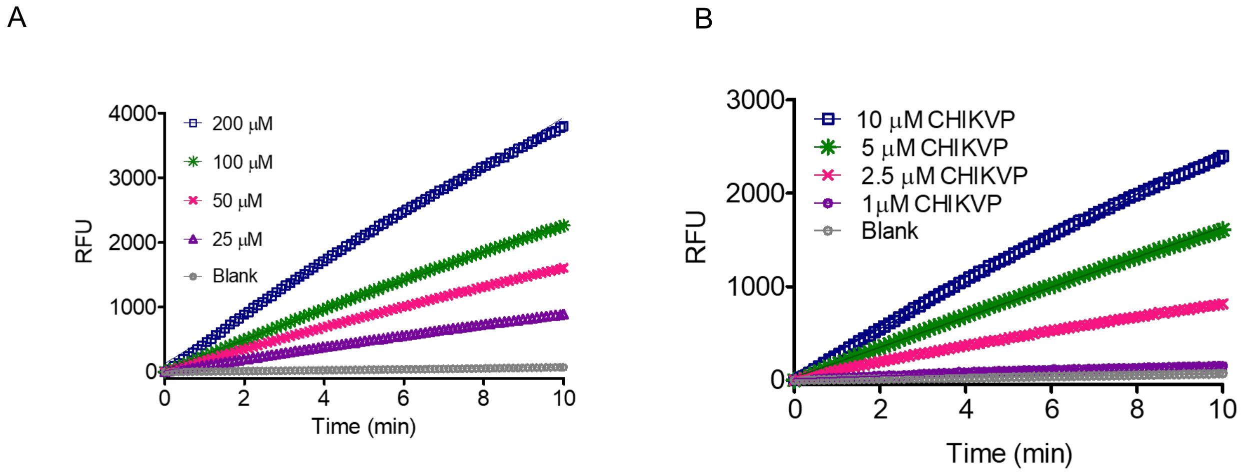


**Figure S9.** Fluorescence based activity assay for CHIKVP with acc-CHIK$_{15}$-dnp (A) with different concentrations of acc-CHIK$_{15}$-dnp substrate (B) with different concentrations of CHIKVP enzyme.

A

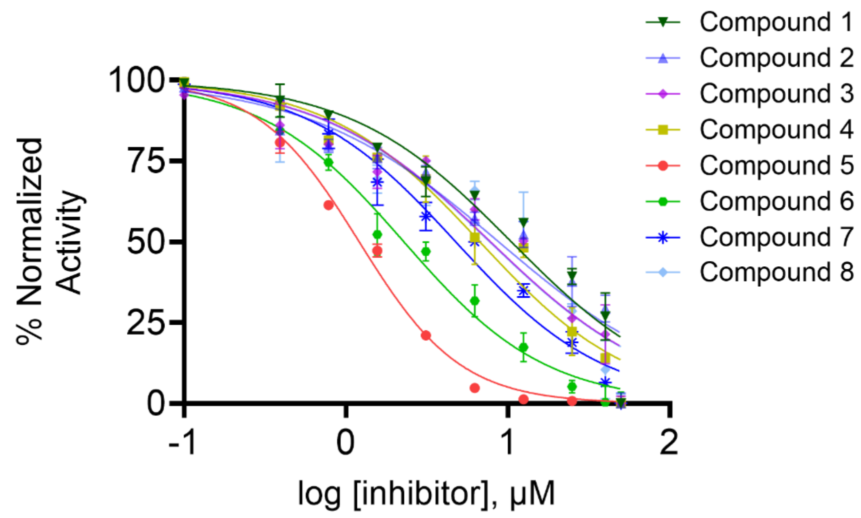


B

| Compound list | $IC_{50}$ (µM) |
|---|---|
| Compound 1 | 10.51 |
| Compound 2 | 8.77 |
| Compound 3 | 7.82 |
| Compound 4 | 6.72 |
| Compound 5 | 1.20 |
| Compound 6 | 2.25 |
| Compound 7 | 4.76 |
| Compound 8 | 7.72 |

**Figure S10.** Inhibition of hit compounds obtained from high throughput screening (HTS) using acc-CHIK$_{15}$-dnp substrate. (A) $IC_{50}$ curves (B) $IC_{50}$ values of the hits.

A

B

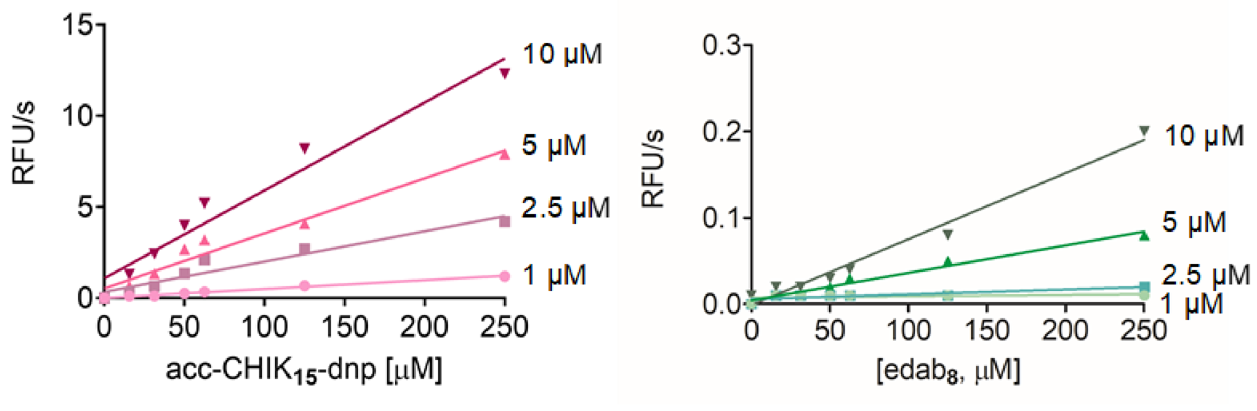


**Figure S11.** Activity of different concentrations of CHIKVP as observed with various concentrations of two peptide substrates. (A) A robust S/N was obtained for lower concentrations of CHIKVP and at lower concentrations of acc-CHIK15-dnp substrate than (B) edab8.

A

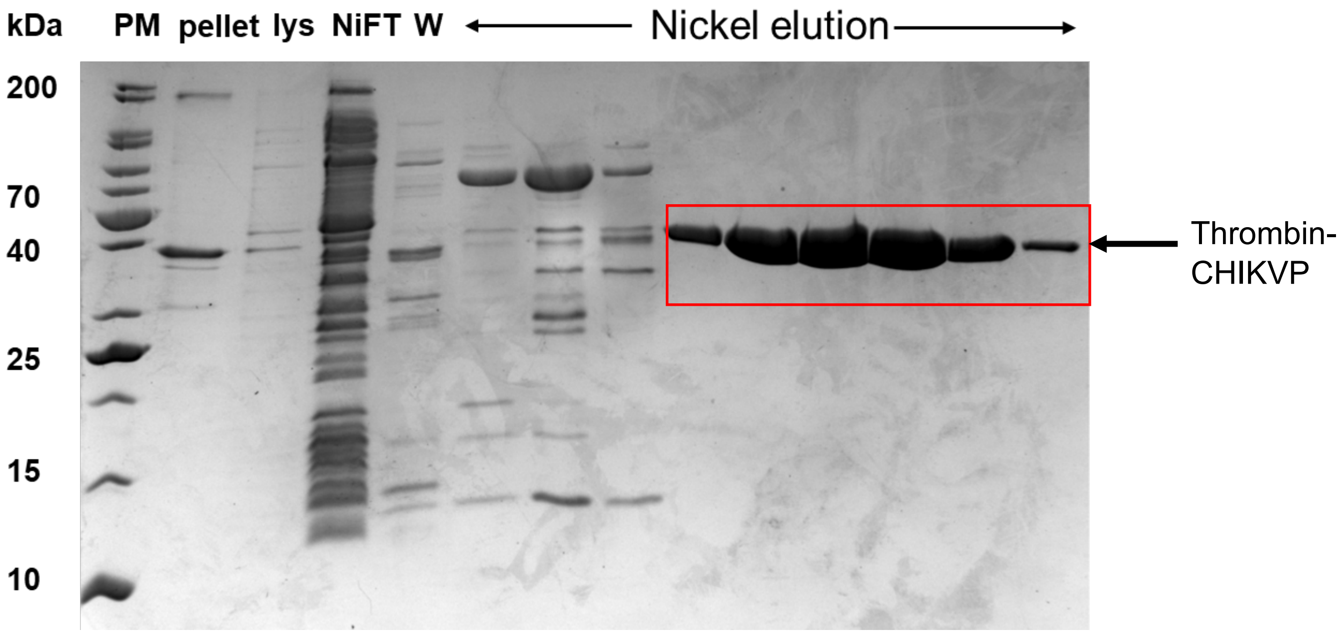

B

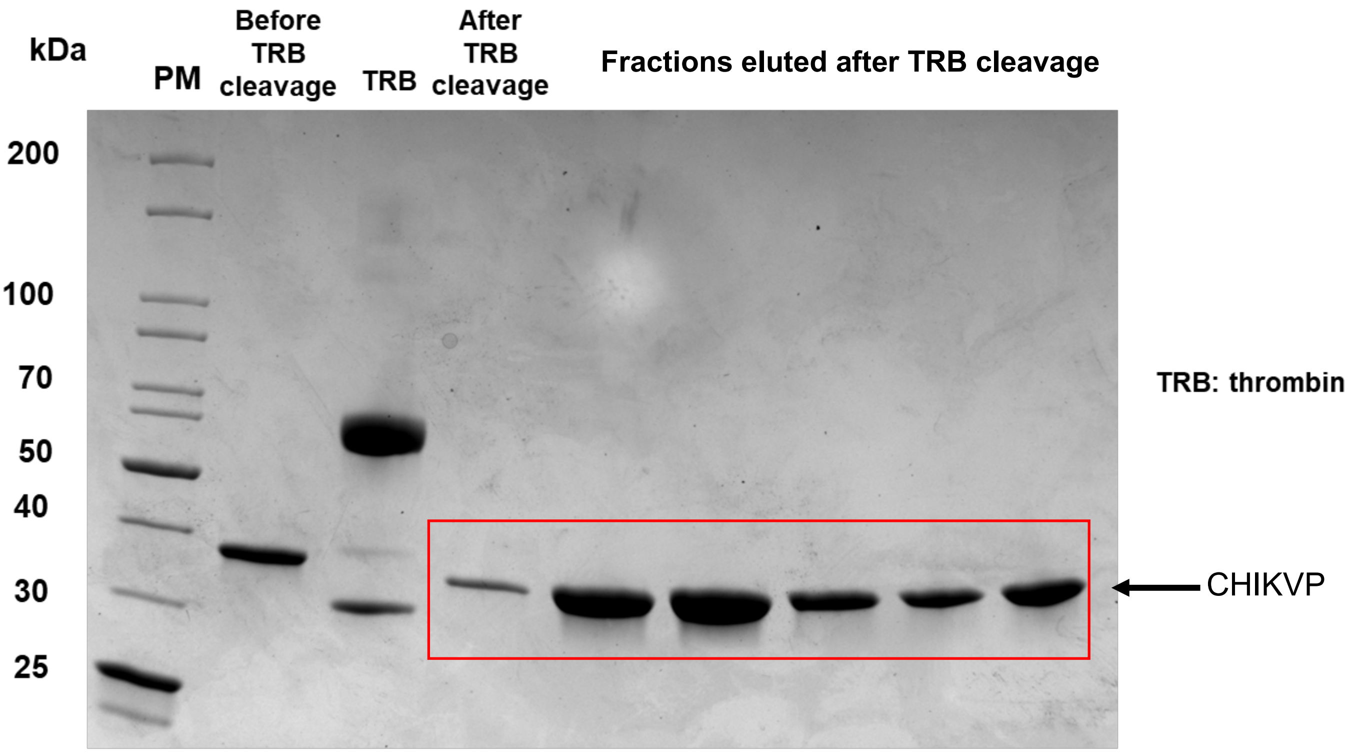


**Figure S12.** Purification of CHIKVP. (A) First nickel column purification. (B) Second nickel column purification after his-tag cleavage.

A

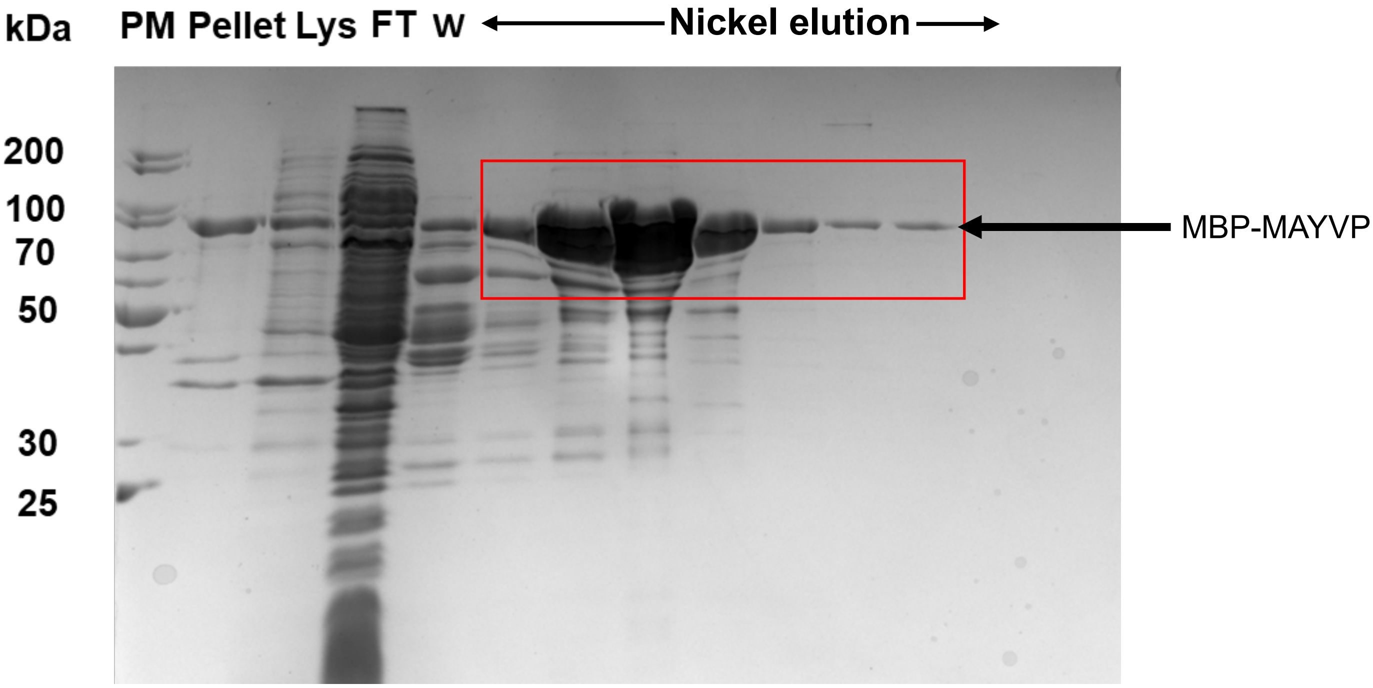

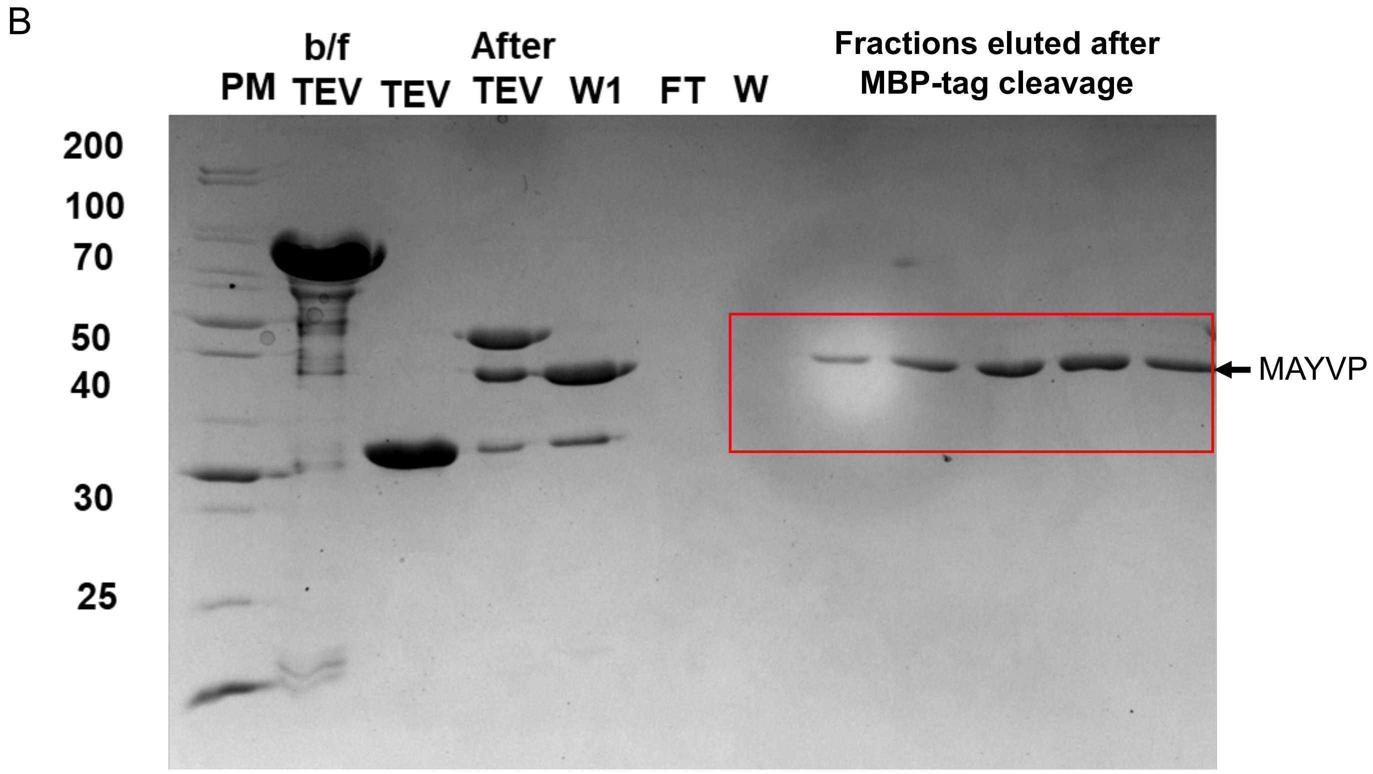


**Figure S13.** Purification of MAYVP. (A) Nickel column purification. (B) MBP column purification after MBP-tag cleavage.

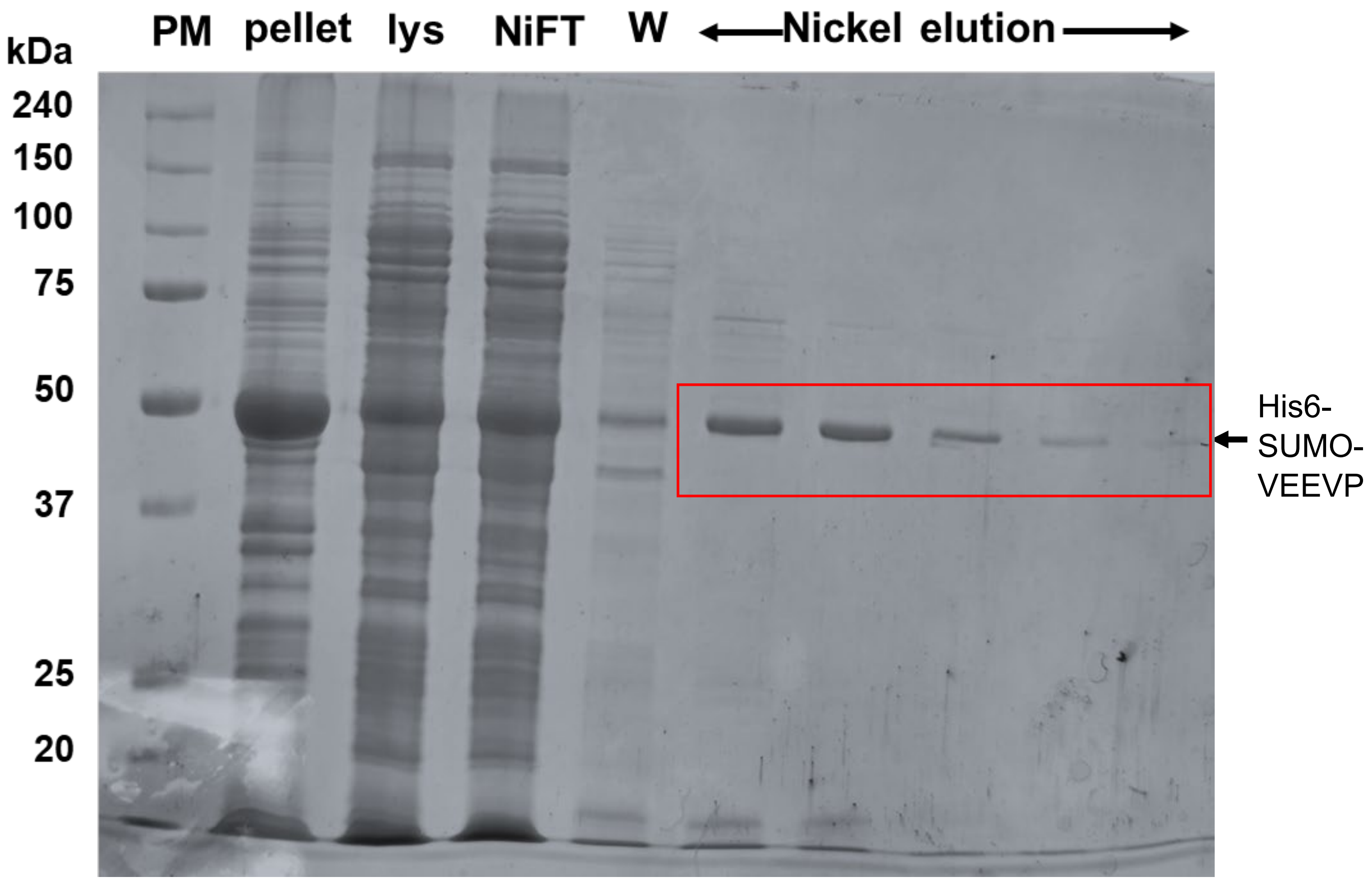


**Figure S14.** Nickel column purification of VEEVP.

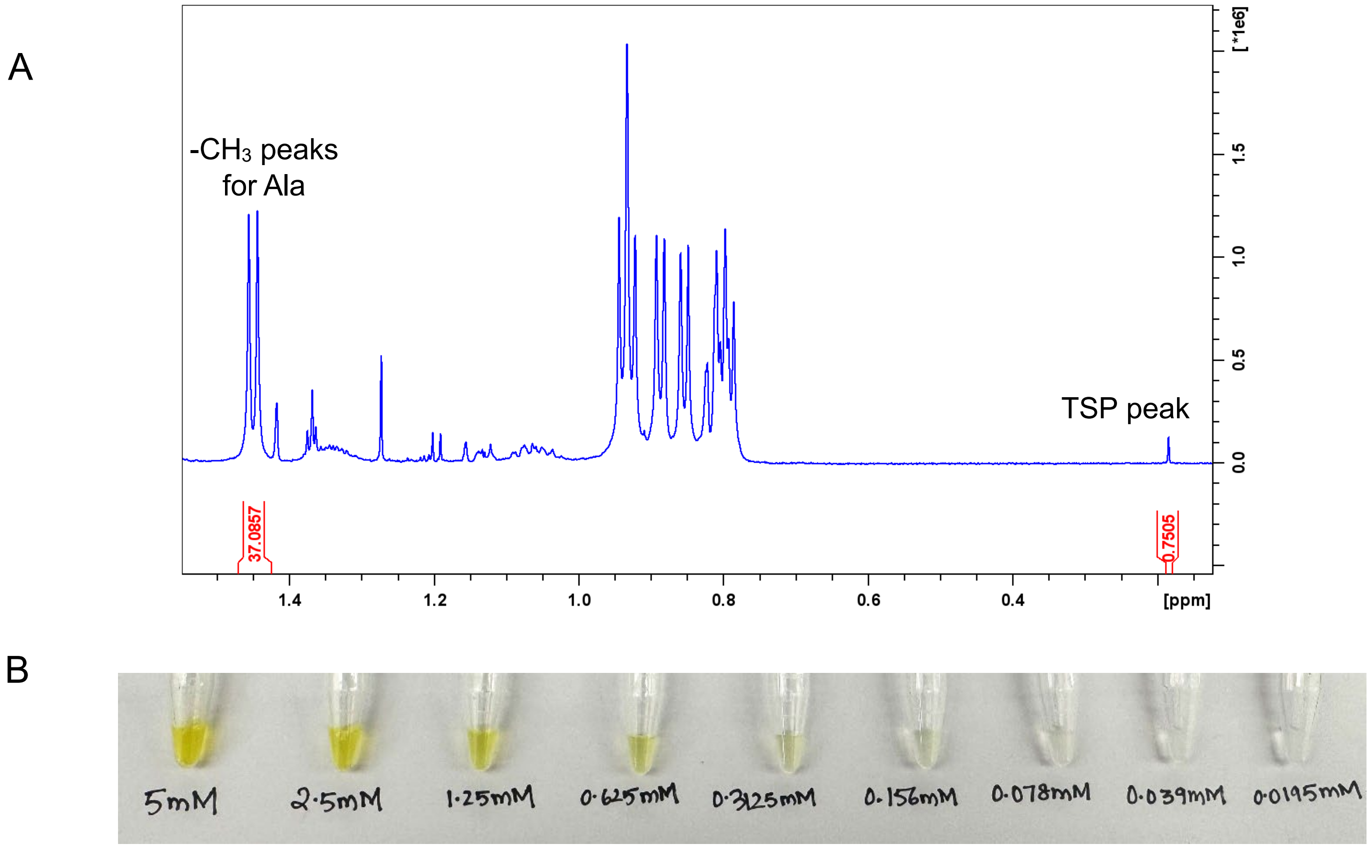


**Figure S15.** Solubility assessment of acc-CHIK$_{15}$-dnp substrate in CHIKVP activity buffer (50 mM Tris pH 7.5, 1mM CHAPS) (A) using $^1$H-NMR at 500 μM which is 10-fold greater concentration than used in the activity assays (50 μM). Trimethylsilyl propionic acid was used as an internal standard. The methyl peaks of alanine were integrated to obtain the concentration of the substrate. The concentration obtained based on the internal standard peak integration was 498 μM, which also excluded the possibility of aggregate formation. (B) No precipitation was observed with naked eye when the substrate was diluted from 5 mM to ten different concentrations as shown.

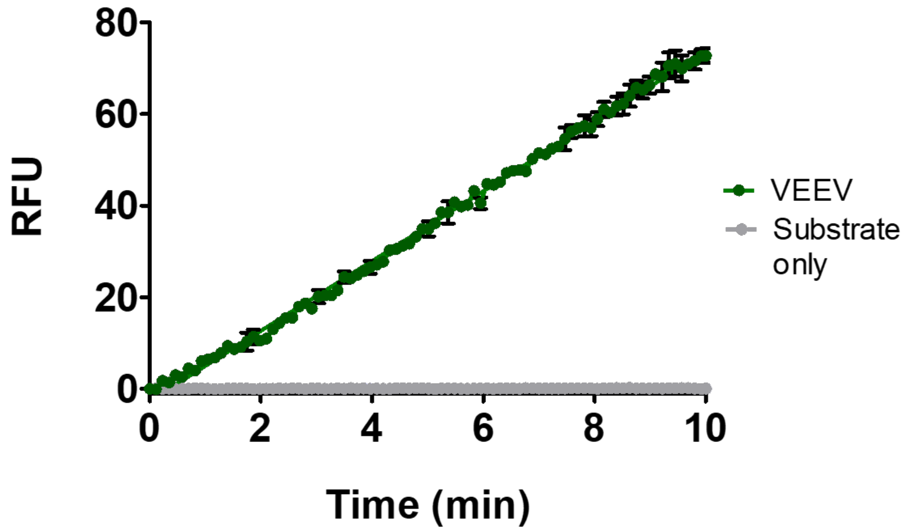


**Figure S16.** Fluorescence based activity assay for 5 μM VEEVP with 25 μM TF5- C-LQEAGA/GSVE-K-(TQ5)-M substrate in 1XPBS, 0.01% triton and 5mM DTT buffer at RT.
